# Neuropilin-1 is a co-receptor for Nerve Growth Factor-evoked pain

**DOI:** 10.1101/2023.12.06.570398

**Authors:** Chloe J. Peach, Raquel Tonello, Elisa Damo, Kimberly Gomez, Aida Calderon-Rivera, Renato Bruni, Harsh Bansia, Laura Maile, Ana-Maria Manu, Hyunggu Hahn, Alex R.B. Thomsen, Brian L. Schmidt, Steve Davidson, Amedee des Georges, Rajesh Khanna, Nigel W. Bunnett

## Abstract

Nerve growth factor (NGF) monoclonal antibodies inhibit chronic pain yet, failed to gain approval due to worsened joint damage in osteoarthritis patients. We report that neuropilin-1 (NRP1) is a co-receptor for NGF and tropomyosin-related kinase A (TrkA) pain signaling. NRP1 is coexpressed with TrkA in human and mouse nociceptors. NRP1 inhibitors suppress NGF-stimulated excitation of human and mouse nociceptors and NGF-evoked nociception in mice. NRP1 knockdown inhibits NGF/TrkA signaling, whereas NRP1 overexpression enhances signaling. NGF binds NRP1 with high affinity and interacts with and chaperones TrkA from the biosynthetic pathway to the plasma membrane and endosomes, enhancing TrkA signaling. Molecular modeling suggests that C-terminal R/KXXR/K NGF motif interacts with extracellular “b” NRP1 domain within a plasma membrane NGF/TrkA/NRP1 of 2:2:2 stoichiometry. G Alpha Interacting Protein C-terminus 1 (GIPC1) scaffolds NRP1 and TrkA to myosin VI and colocalizes in nociceptors with NRP1/TrkA. GIPC1 knockdown abrogates NGF-evoked excitation of nociceptors and pain-like behavior. NRP1 is a nociceptor-enriched co-receptor that facilitates NGF/TrkA pain signaling. NRP binds NGF and chaperones TrkA to the plasma membrane and signaling endosomes via the GIPC1 adaptor. NRP1 and GIPC1 antagonism in nociceptors offers a long-awaited non-opioid alternative to systemic antibody NGF sequestration for the treatment of chronic pain.

**Summary:** Neuropilin-1 and G Alpha Interacting Protein C-terminus 1 are necessary for nerve growth factor-evoked pain and are non-opioid therapeutic targets for chronic pain.

## Introduction

The first discovered growth factor, nerve growth factor (NGF), was identified by its ability to stimulate growth of sympathetic neurons (1). Tropomyosin-related kinase A (TrkA), a receptor tyrosine kinase (RTK), mediates these neurotrophic actions of NGF (2). After NGF binds to TrkA at peripheral nerve terminals, stable NGF/TrkA signalosomes are transported in a retrograde direction to the distant neuronal soma, where they regulate transcription and neuronal development (3, 4). NGF and TrkA are also strongly implicated in multiple forms of pain (5). p75NTR is another receptor for NGF, as well as the immature pro­ NGF peptide (6, 7) that activates opposing pro-apoptotic signaling pathways (8). Although NGF and its receptors have been intensively studied in the context of neuronal development and pain, mechanistic gaps in understanding have hampered the successful approval of NGF-directed therapeutics.

Chronic pain afflicts twenty percent of the global population at some point in life, yet is inadequately treated by non-steroidal anti-inflammatory drugs and opioids, which lack efficacy and have life-threatening side effects. NGF/TrkA is one of the few non-opioid targets for chronic pain validated in patients. NGF, which is produced by injured and diseased tissues, activates TrkA on peripheral nociceptors to evoke rapid sensitization and stimulate the transcription of neuropeptides, receptors and ion channels that mediate pain (5). NGF-stimulated neuronal sprouting may also contribute to pain and accompanies painful conditions (5). NGF and TrkA have been implicated in pain associated with inflammation, nerve injury and cancer (5). The central role of TrkA and NGF in pain is evident in patients with Hereditary Sensory and Autonomic Neuropathy (HSAN) Type IV and V, where pathological insensitivity to pain results from loss-of-function mutations in TrkA (9) and NGF (10), respectively. Although monoclonal antibodies (mAbs) are analgesic in osteoarthritic patients (11), they lack FDA approval due to worsening joint damage in some individuals (12). The identification of nociceptor-enriched mediators of NGF-induced pain may facilitate development of non­ opioid analgesics, avoiding the adverse effects of systemic NGF sequestration with mAbs.

Transcriptome analyses have identified proteins that are conserved between rodent and human nociceptors, including neuropilin-1 (NRP1) (13). NRP1 is a type I transmembrane protein discovered for its role in axon guidance (14). Lacking a catalytic domain, NRP1 does not transduce signals *per se* but rather acts as a co-receptor for unrelated families of proteins, including vascular endothelial growth factor (VEGF-A) (15). NRP1 is a co-receptor with VEGF receptor 2 (VEGFR2) that enhances VEGF-A-induced blood vessel development. NRP1 overexpression in cancers spurred the development of a mAb, Vesencumab/MNRP1685A, to inhibit tumorigenesis (16). NRP1 is also implicated in VEGF-A-mediated pain (17, 18). Since NGF, a major pain mediator, possesses a prospective NRP1 binding motif, we investigated the hypothesis that NRP1 is a co-receptor for NGF and TrkA in nociception.

## Results

### NRP1 binds human -NGF

NRP1 is a co-receptor for unrelated families of proteins including neuronal guidance molecules (semaphorins) (19), growth factors (VEGF-A) (15), cell-penetrating peptides (20), and viruses (SARS-CoV-2) (14). NRP1 ligands share a C-terminal basic motif (C-end rule, “CendR” motif, R/KXXR/K), which interacts with extracellular “b” domains of NRP1 (15, 19). Inspection of the amino acid sequence of mature NGF identified prospective ‘CendR’ R/KxxR/K motifs within the C-terminus that are conserved between rodents and humans and known to mediate the interaction of other growth factors with NRP1 (20) (Figure 1A).

**Figure 1.**
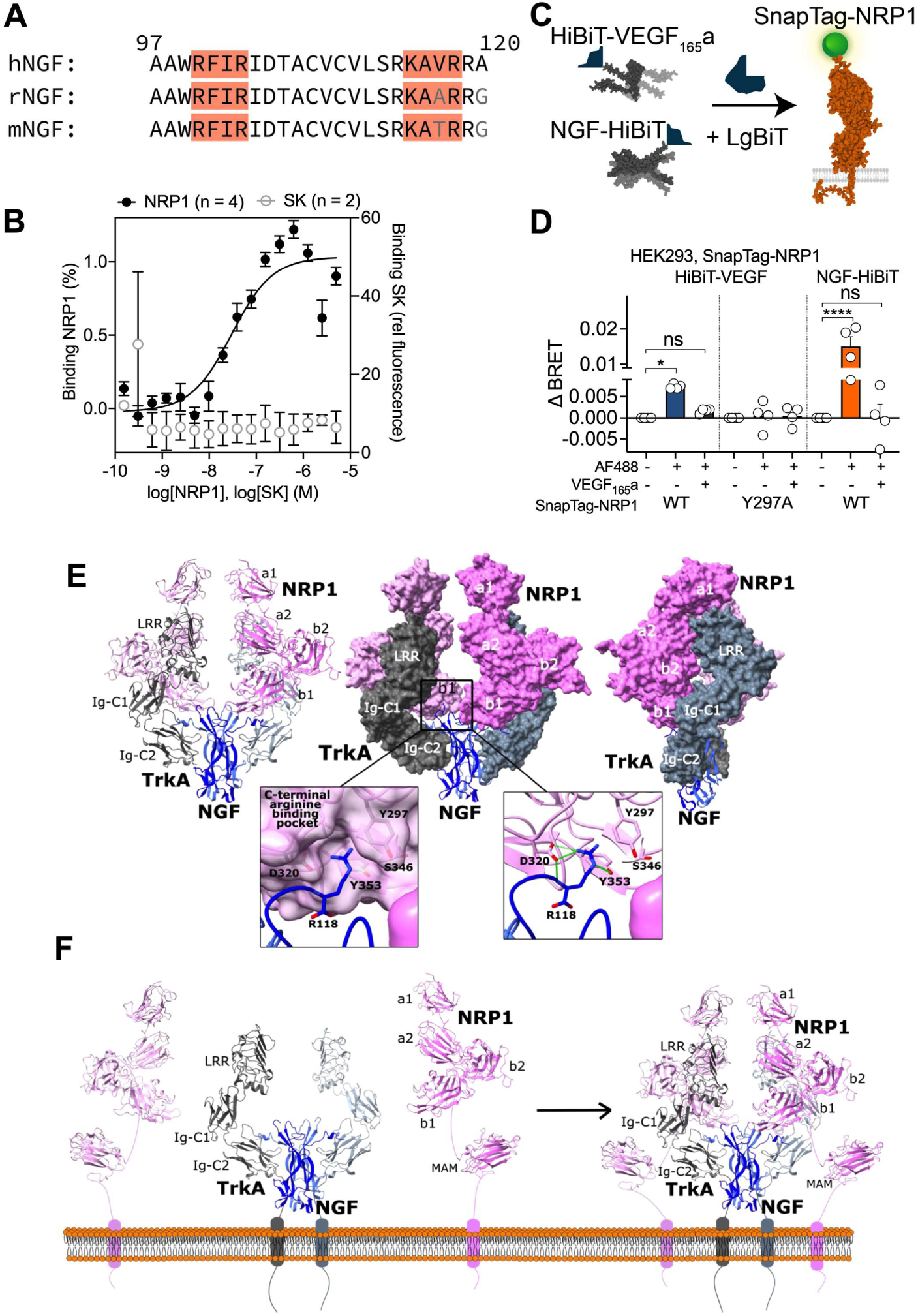
NGF interacts with NRP1. **A.** Carboxy-terminal sequences of NGF, highlighting prospective CendR motifs (R/KxxR/K). Amino acids numbered according to the mature NGF sequence (1-120 equivalent to proNGF 122-241). *h, Homo sapiens; r, Rattus norvegicus; m, Mus musculus.* **B.** Interaction between fluorescent human NGF and human NRP1 (residues 22-644) or staphylokinase (SK, negative control) and using MST. Data from 4 (NRP1) or 2 (SK) independent experiments. Mean±SEM. **C, D.** BRET measurements ofVEGF and NGF proximity (<10 nm) to full-length NRP1 in living HEK293T cells at 37°C **(C).** Supernatant was collected from cells secreting growth factor (VEGF_165_a or NGF) tagged with HiBiT, a small fragment of nanoluciferase. When reconstituted with recombinant LgBiT and luciferase substrate, this acts as a bioluminescent donor for SnapTag-NRP1 labeled with SNAPTag-Alexa Fluor® 488 (AF488). BRET was compared to negative control (HiBiT/LgBiT only lacking AF488). Cells were pre-incubated with vehicle or 10 nM unlabeled VEGF15sa (30 min), followed by luminescent growth factor (15 min, 37°C). BRET with HiBiT-VEGF_165_a was measured between HiBiT-VEGF_165_a and either SnapTag-NRP1 (WT) or the known VEGF_165_a binding-dead mutant (Y297A), as well as HiBiT-tagged NGF (D). Data from 4 independent experiments with triplicate wells. Mean±SEM. *P<0.05, ****P<0.0001, ns not significant. 1-way ANOVA, Sfdak’s multiple comparisons. **E, F.** Ternary complex of human NGF/TrkA/NRP1 generated using a data constraint-driven computational docking protocol. Cartoon and surface representation of TrkA (grey shades), NGF (blue shades) and NRP1 (pink shades) are shown. **E.** NGF/TrkA/NRP1 docked model and conserved molecular interactions at the NGF/NRP1 interface. The docked model of NGF/TrkA/NRP1 suggests a 2:2:2 stoichiometry with one NRP1 molecule interacting with one TrkA molecule and the NGF dimer. Two views of the docked complex with NRP1 and TrkA in surface representation are shown to represent shape complementarity between NRP1 and TrkA. Binding of NGF C-terminal R118 (blue sticks) to conserved residues (pink sticks, Y297, D320, S346, Y353) in C-terminal arginine binding pocket located in NRP1 b1 domain is shown as inset with predicted hydrogen bonding interactions depicted as green dashes. **F.** Illustration of the proposed complex formation between NRP1 and NGF/TrkA as it would appear on the cell surface. The membrane proximal MAM domains of NRP1 (PDB 5L73) are included for completeness (MAM domains were not part of docking calculations) to propose a sterically feasible membrane tethered NGF/TrkA/NRP1 complex formation on the cell surface. The membrane linkers connecting NRP1 b2 and MAM domains, linkers connecting NRP1, TrkA with their respective transmembrane regions and the transmembrane regions are not derived from structured components and are not to scale.

To directly investigate whether NGF binds NRP1, microscale thermophoresis (MST) was used to analyze interactions between fluorescent human NGF and the extracellular a1a2b1b2 domain of human NRP1. NGF interacted with NRP1 with low nanomolar affinity (Kd = 35.5 ± 4.7 nM, N=4, Figure 1B). No interaction was detected between fluorescent NGF and staphylokinase (negative control). The measured NGF/NRP1 affinity is lower than the reported affinities of NGF for TrkA and p75NTR, which are in the 1-15 nM range using surface plasmon resonance (21, 22), but is comparable to affinity of VEGF_165_a for NRP1 (Kd 9 nM (23) to 120 nM (24)).

NGF binding to NRP1 in living cells at 37°C was analyzed using a modified bioluminescence resonance energy transfer (BRET)-based assay (Figure 1C). VEGF15sa (positive control growth factor known to interact with NRP1) or NGF was genetically fused to an 11 amino acid fragment (HiBiT) of nanoluciferase (Nanoluc). HiBiT has a high affinity for the complementary LgBiT fragment. HiBiT-tagged growth factors were expressed in HEK293T cells, secreted into supernatant, and conjugated to recombinant LgBiT to form full length and catalytically active Nanoluc. Upon addition of furimazine substrate, complemented Nanoluc emits bioluminescence. This supernatant containing HiBiT-tagged growth factor was incubated with cells expressing SnapTag-NRP1 labeled with SNAP-Surface® Alexa Fluor® 488 fluorophore (SnapTag-AF488, +) or lacking fluorophore (-). If proteins are in close proximity (<10 nm), luminescent growth factor can act as a BRET donor to fluorescent NRP1. As a positive control, BRET was detected between luminescent VEGF_165_a and SnapTag-NRP1 (Figure 1D). Pre-incubation of cells with an excess of VEGF,s5a (10 nM) abolished the BRET signal. No BRET signal was detected when luminescent VEGF155a was incubated with cells expressing a binding-dead NRP1 mutant in the b1 domain (25, 26). Importantly, BRET was detected between NGF-HiBiT and SnapTag-NRP1, and pre-incubation with unlabeled VEGF155a inhibited this response.

The results from analysis of interactions between recombinant proteins and ligand binding studies in intact cells provide evidence that NGF directly binds NRP1 with nanomolar affinity.

### Modeling an NGF/TrkA/NRP1 ternary complex

Ternary complexes of human NGF/TrkA/NRP1 were generated through an information-driven computational docking protocol using HADDOCK 2.4 (27). The complexes were analyzed against available biochemical data. Binding interactions of sterically feasible complexes in the cellular context were then analyzed at the molecular level. NRP1 is composed of five extracellular domains from N- to C-terminus, namely a1, a2, b1, b2 and c/MAM (28). The extracellular domain (ECO) of TrkA comprises an N-terminal leucine-rich repeat (LRR) domain and C-terminal immunoglobulin domains (lg-C1, lg-C2) (29). The crystal structure of NGF/TrkA complex reveals a 2:2 stoichiometry (PDB 2IFG). Our docked model of NGF/TrkA/NRP1 suggests a 2:2:2 stoichiometry with one NRP1 molecule interacting with one TrkA molecule and the NGF dimer (Figure 1E, Supplemental Table S1). The model suggests that the NRP1 b1 domain predominantly interacts with the NGF C-terminus while also interfacing with an interdomain region between lg-C1 and lg-C2 domains of TrkA. In this model, the NRP1 a1 and a2 domains interface with the top of the TrkA LRR domain and the NRP1 b2 domain slots in the curved space between TrkA lg-C1 and LRR domains contributing to shape complementarity between TrkA and NRP1 (Figure 1E). NGF has two prospective ‘CendR’ R/KxxR/K motifs within the C-terminus that mediate interaction of other growth factors with NRP1 b1 domains (20, 30, 31). We observed that in this model, R118 from CendR motif K115AVR11B binds to C-terminal arginine binding pocket located in the NRP1 b1 domain through molecular interactions, including hydrogen bonding interactions between NGF R118 and NRP1 conserved residues Y297, D320 and Y353 (Figure 1E inset). NRP-binding proteins and peptides possess a C-terminal arginine that binds to a specific C-terminal arginine binding pocket of the NRP1 b1 domain through similar set of molecular interactions (31, 32). The membrane proximal MAM domain of NRP1, extending from the NRP1 b2 domains, is proposed to orient the other extracellular domains of NRP1 away from the membrane for protein/protein interaction (28). In the NGF/TrkNNRP1 docked model, the NRP1 b2 domain lies perpendicular to the NGF/TrkA 2-fold symmetry axis with solvent accessible C-termini pointing towards the membrane (Figure 1E). This orientation suggests that it is possible to extend the MAM domain from the b2 domain towards the membrane while NRP1 is still bound to the NGF/TrkA complex resulting in a sterically permissible membrane tethered NGF/TrkA/NRP1 complex (Figure 1F). TrkA Asn-linked glycosylation can regulate its localization and activity (33). Although non-standard residues were excluded from docking calculations, the NGF/TrkNNRP1 docked model shows that NRP1 binding to TrkA does not clash with Asn­ linked glycosylations on TrkA suggesting sterically feasible binding of NRP1 to glycosylated TrkA in a cellular context (Supplemental Figure S1). Thus, our analysis provides structural insights into the NRP1 co­ receptor function for NGF/TrkA signaling analogous to a recent structural study of NRP1 co-receptor function with an unrelated family of proteins (34).

### *Ntrk1/TrkA* and Nrp1/NRP1 are coexpressed in mouse and human DRG neurons

To determine whether NRP1 could function as an NGF/TrkA co-receptor in nociceptors, TrkA and NRP1 were localized in mouse and human dorsal root ganglia (DRG) by immunofluorescence (protein) and RNAScope™ *in situ* hybridization (mRNA). In mice, immunoreactive TrkA was detected in vesicles and immunoreactive NRP1 was localized at the plasma membrane of the same DRG neurons (Figure 2A). *Ntrk1* (TrkA) and *Nrp1* (NRP1) mRNAs also colocalized in mouse DRG neurons, which were identified by NeuN immunostaining (Figure 2B). *Nrp1* was detected in 42% of small-diameter peptidergic nociceptors of mouse DRG that expressed immunoreactive calcitonin-gene-related peptide (CGRP) (Figure 2C). lmmunoreactive NRP1 was also detected in satellite glial cells of DRG, which were identified by immunostaining for glutamine synthetase (GS) (Figure 2D). In humans, *Ntrk1* and *Nrp1* mRNAs also colocalized in DRG neurons (Figure 2E). *Nrp1* was detected in 59% of human DRG that expressed immunoreactive CGRP and 42% of neurons expressing P2X purinoceptor 3 (P2X3) (Figure 2F). In mice, *Nrp1* mRNA was expressed in 33% of Ntrk1-positive neurons (Figure 2G). In humans, *Nrp1* was coexpressed in 93% of *Ntrk1-* positive neurons (Figure 2H). Thus, TrkA and NRP1 are coexpressed in the same neurons in DRG in mouse and human, where NRP1 is appropriately located to control NGF/TrkA signaling.

**Figure 2.**
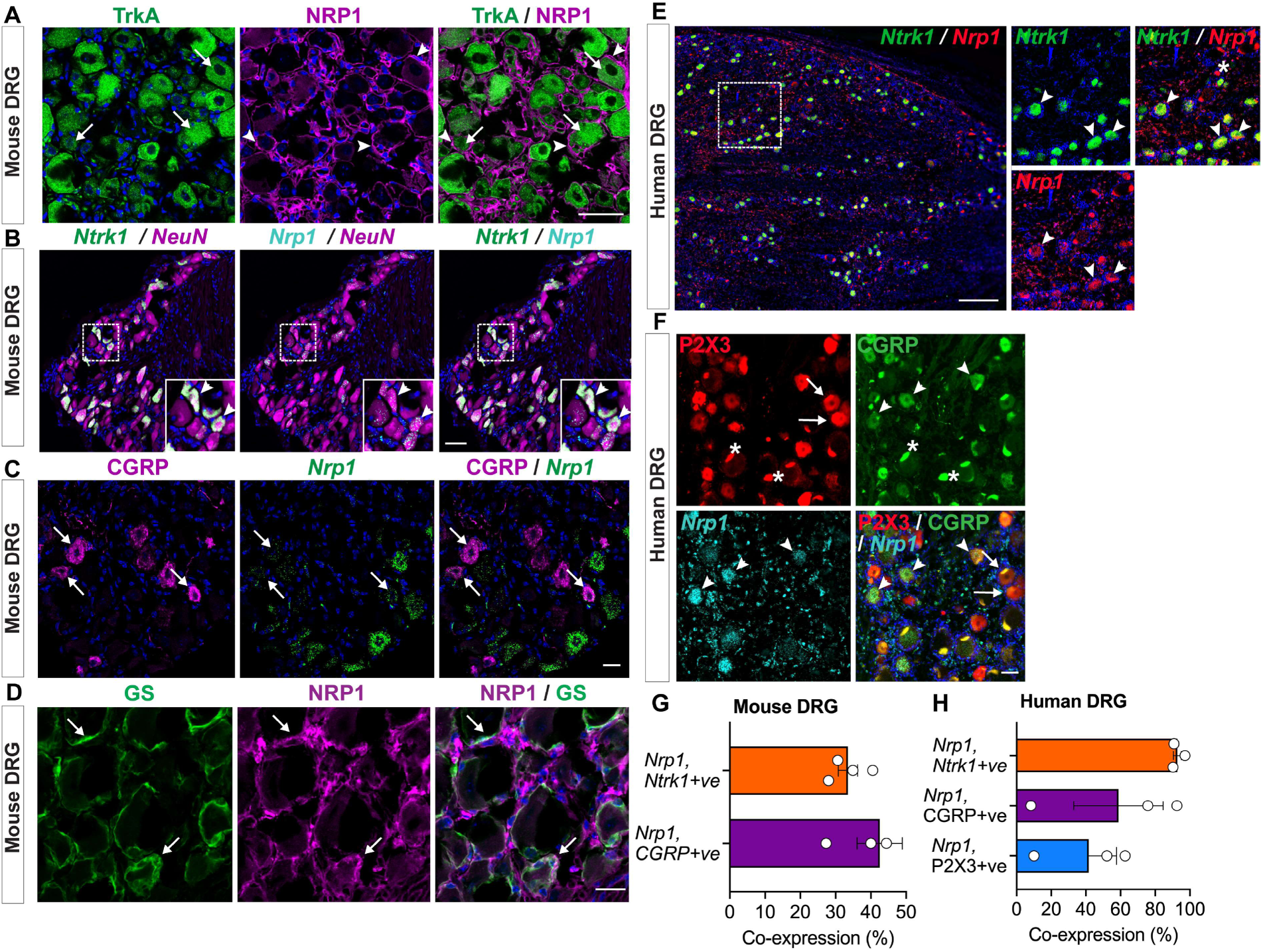
TrkA and NRP1 are coexpressed in DRG. **A.** lmmunofluorescence detection of TrkA and NRP1 in mouse DRG. TrkA was largely intracellular (arrows) whereas NRP1 was localized to the plasma membrane (arrow heads). Scale bar, 50 µm. **B.** RNAScope™ detection of *Ntrk1* (TrkA) and *Nrp1* (NRP1) mRNA in mouse DRG neurons identified by NeuN immunofluorescence. Arrow heads indicate neurons coexpressing *Ntrk1* and *Nrp1.* Scale bar, 50 µm. **C.** lmmunofluorescence detection of CGRP and RNAScope™ detection of *Nrp1* mRNA in mouse DRG. Arrows indicate neurons coexpressing CGRP and *Nrp1.* Scale bar, 20 µm. **D.** lmmunofluorescence detection of NRP1 and glutamine synthetase (GS) in mouse DRG. Arrows indicate satellite glial cells expressing NRP1. Scale bar, 50 µm. **E.** RNAScope^TM^ detection of *Ntrk1* and *Nrp1* mRNA in human DRG. Arrow heads indicate neurons coexpressing *Ntrk1* and *Nrp1.* Scale bar, 500 µm. **F.** lmmunofluorescence of P2X3 and CGRP and RNAScope™ detection of *Nrp1* mRNA in human DRG. Arrow heads indicate neurons coexpressing CGRP and *Nrp1.* Arrows indicate neurons expressing P2X3 but not *Nrp1.* Scale bar, 50 µm. *Denotes fluorescence in human neurons due to lipofuscin. Nuclei shown in blue. **G.** Percentage of mouse DRG neurons expressing *Ntrk1* or CGRP that coexpress *Nrp1.* **H.** Percentage of human DRG neurons expressing *Ntrk1,* CGRP or P2X3 that coexpress *Nrp1.* A-F show representative images from N=4-5 mice and N=3 humans. G, H show hybridized positive neurons(%) from N=3-4 mice and N=3 humans.

### NRP1 inhibitors suppress NGF-induced sensitization of TRPV1 in mouse DRG neurons

The transient receptor potential vanilloid-1 (TRPV1) ion channel mediates the algesic actions of diverse painful stimuli and is sensitized by many receptors, including TrkA (5, 35). The contribution of NRP1 to NGF-induced sensitization of TRPV1 was determined by calcium imaging of mouse DRG neurons. Exposure of neurons to the TRPV1 agonist capsaicin (100 nM) increased [Ca^2^+]i by 239.3 ± 137.3% (mean ± SD, N=1696 cells) of basal, consistent with TRPV1 activation (Figure 3A). The response to a second capsaicin challenge 6 min later was reduced to 96.13 ± 23.60% (N=1696 cells) of the first response, indicating TRPV1 desensitization (P<0.001, paired t-test). When neurons were incubated with mouse NGF (100 nM) for 2 min before the second challenge, the response to the second capsaicin challenge was amplified to 119.3 ± 45.26% (N=1621 cells) of the first response, denoting TRPV1 sensitization (P<0.001, paired t-test).

**Figure 3.**
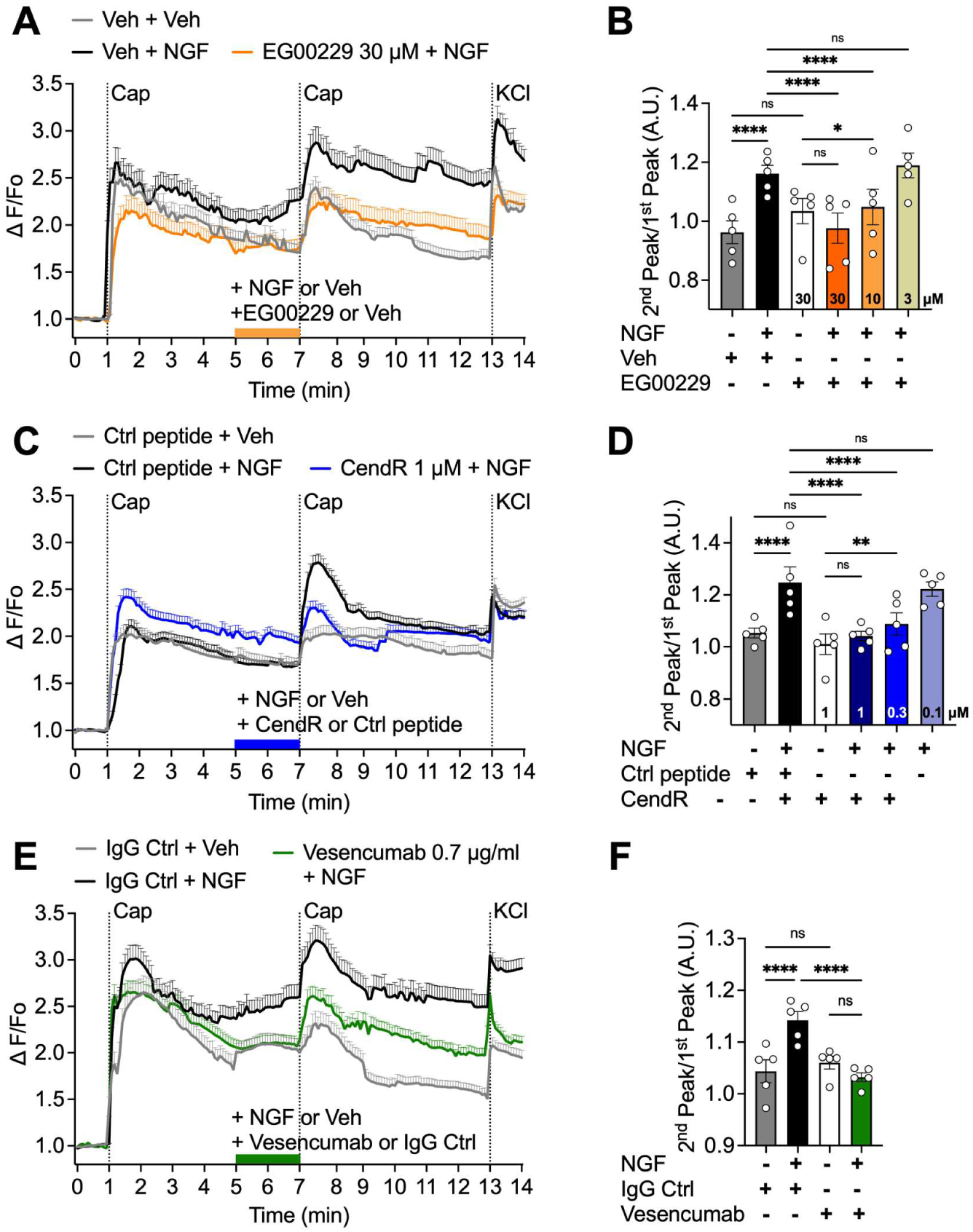
NRP1 inhibition prevents NGF-induced sensitization of TRPV1. **A, C, E.** Time course of responses of mouse DRG neurons to repeated challenge with capsaicin (Cap, 100 nM) expressed as ΔF/Fo ratio. **B, D, F.** Summary of responses to capsaicin expressed as the ΔF/Fo of the second capsaicin Ca^2+^ response over the ΔF/Fo of the first capsaicin Ca^2+^ response (2^nd^ peak/1^st^ peak). **A, B.** Effect of NRP1 inhibitor EG00229 (3, 10, 30 µM, 30 min pre-incubation) or vehicle (Veh). **A.** N=64-119 cells per trace. **B.** Summary from N=5 independent experiments. **C, D.** Effect of NRP1 inhibitor CendR or control (Ctrl) (0.1, 0.3, 1 µM). **C.** N=142-535 cells per trace. **D.** Summary from N=5 independent experiments. **E, F.** Effect of a human mAb against the b1b2 domain of NRP1 (Vesencumab) or control (Ctrl) lgG (0.7 µg/ml). **E.** N=97-199 cells per trace. **F.** Summary from N=5 independent experiments. A.U., arbitrary units. Mean±SEM. *P<0.05, **P<0.01, ****P<0.0001, ns not significant. Two-way ANOVA with Tukey’s multiple comparisons test.

The contribution of NRP1 to NGF-induced sensitization of TRPV1 was examined by incubating DRG neurons with NRP1 antagonists or control reagents before the second capsaicin challenge. EG00229 is a small molecule inhibitor developed to inhibit binding of VEGF-A to the b1 domain of NRP1 (25). EG00229 (3, 10, 30 µM) had concentration-dependent effects on NGF-induced sensitization of TRPV1 compared to vehicle control (0.1% DMSO) (Figure 3A, B). EG00229 30 µM completely prevented NGF­ induced sensitization of TRPV1, 10 µM EG00229 partially inhibited TRPV1 sensitization, and 3 µM EG00229 was ineffective. To competitively inhibit binding of NGF to NRP1, a peptide fragment of NGF was synthesized that includes the two conserved ‘CendR’ R/KxxR/K motifs within the C-terminus of NGF (underlined, QAAWR<u>FIR</u>IDTACVCVLSR<u>KAVR</u>RA, corresponds to position 96-120 of mature NGF peptide), which were predicted by molecular modeling to interact with the b1 domain of NRP1. A peptide fragment of NGF that was not predicted to interact with NRP1 (ARVAGQTRNITVDPRLFKKRRLRSP, corresponds to position 61-85 of immature NGF peptide) was used as a control (Ctrl peptide). The CendR peptide (0.1, 0.3, 1 µM) also had concentration-dependent effects on NGF-induced sensitization of TRPV1 compared to a Ctrl peptide. CendR at 0.3 or 1 µM prevented TRPV1 sensitization whereas 0.1 µM CendR was ineffective (Figure 3C, D). Vesencumab, a human mAb against the b1b2 domain of NRP1 (16) (0.7 µg/ml), similarly prevented NGF-induced sensitization of TRPV1 compared to control lgG (Figure 3E, F). None of the inhibitors affected the response to capsaicin in neurons that were not treated with NGF. The finding that three mechanistically distinct inhibitors of NRP1 (EG00229, CendR peptide, Vesencumab mAb) reproducibly blocked NGF-induced sensitization ofTRPV1 suggests that NRP1 controls the pronociceptive actions of NGF and TrkA, potentially by enhancing the signaling competency of the NGF-TrkA complex that sensitizes TRPV1.

### NRP1 inhibitors suppress NGF-induced ionic currents in mouse and human DRG neurons

NGF/TrkA signaling promotes pain by kinase-mediated phosphorylation and sensitization of ion channels, leading to enhanced sensitivity to mechanical, thermal and chemical stimuli, thus activating voltage-gated ion channels in DRG neurons (5). Patch-clamp recordings under current-clamp mode were made to determine whether NRP1 is necessary for NGF-induced excitability and ion channel activity in dissociated DRG neurons. Exposure of mouse DRG neurons to mouse NGF (50 nM) stimulated action potential firing of nociceptors elicited by a depolarizing ramp pulse (Figure 4A, B). Pre-incubation with the NRP1 inhibitor EG00229 at 30 µM, but not 10 µM, prevented NGF-induced hyperexcitability (Figure 4A, B). NGF or EG00229 did not affect the resting membrane potential (Figure 4C) or the rheobase, the minimum current necessary to elicit an action potential (Figure 4D). Since EG00229 at 10 µM did not affect neuronal excitability, we proceeded with testing only the higher concentration (30 µM) in subsequent experiments.

**Figure 4.**
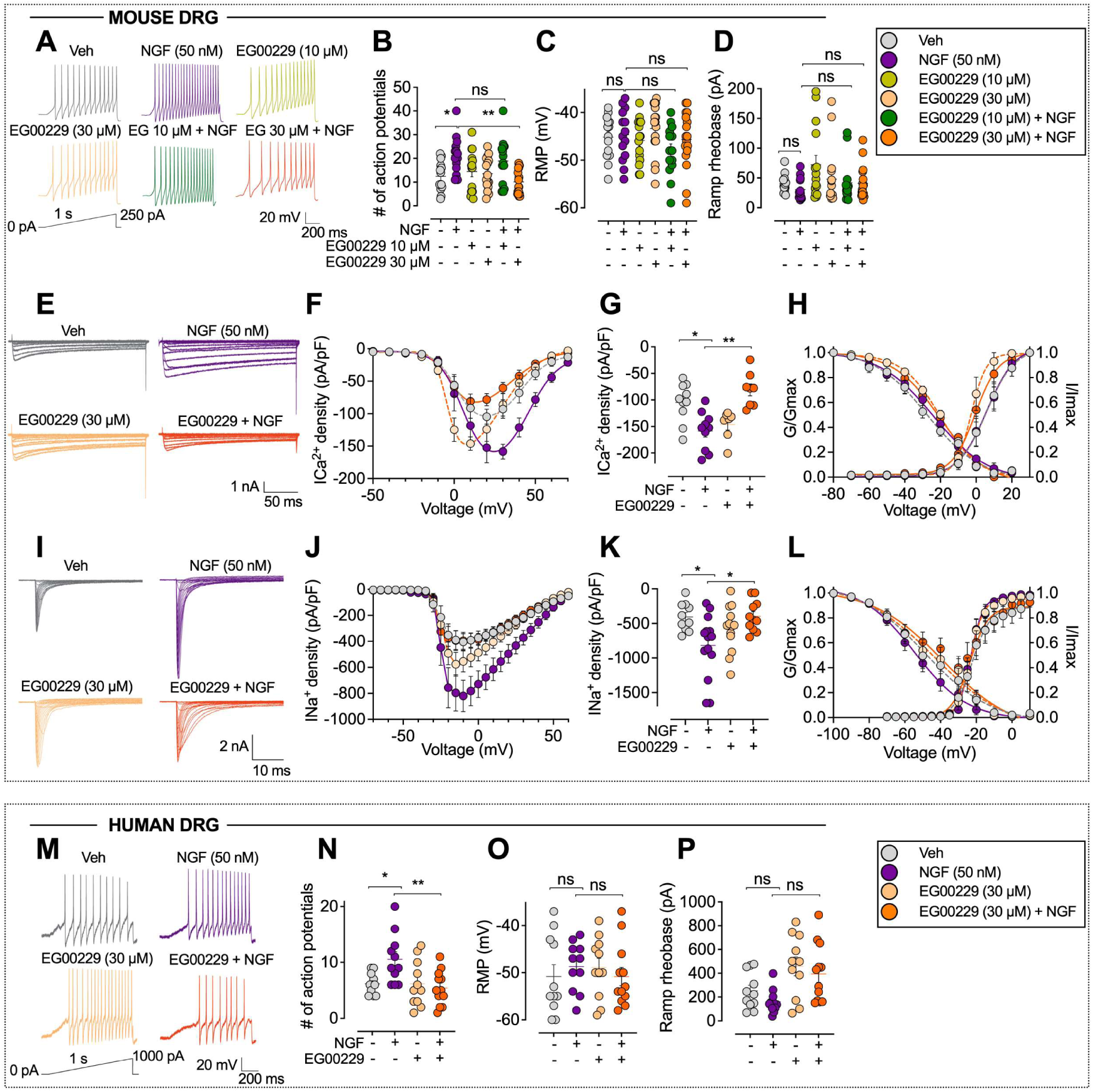
NRP1 inhibition prevents NGF-induced increases in neuronal firing, excitability and ion channel currents. Effect of NRP1 inhibitor EG00229 (10 or 30 µM, 30 min) on responses of dissociated mouse **(A-L)** and human **(M-P)** DRG neurons to NGF (50 nM, 30 min). **A, B, M, N.** Representative action potential firing evoked by a depolarizing ramp stimulus **(A, M),** with summary of the number of evoked action potentials **(B, N).** N=14-17 cells **(B-D),** N= 11-12 cells **(N-P). C, D, 0, P.** Resting membrane potential (RMP) **(C, 0)** and ramp rheobase **(D, P). E-H.** Representative family of Ca^2+^ current traces recorded from small diameter DRG neurons in response to depolarization steps from -70 to +70 mV from a holding potential of -90 mV **(E),** with double Boltzmann fits for current density-voltage curve **(F),** summary of peak calcium current densities **(G)** and Boltzmann fits for voltage-dependence of activation and inactivation **(H).** N=7-10 cells. **I-L.** Representative family of Na^+^ current traces, where currents were evoked by 150 ms pulses between -70 and +60 mV (I), with double Boltzmann fits for current density-voltage curve **(J),** summary of peak sodium current densities **(K)** and Boltzmann fits for voltage-dependence of activation and inactivation **(L).** N=10-14 cells. Mean±SEM. *P<0.05, ** P<0.01, ns not significant. B, C, D, G, K, N, 0, P. Kruskal-Wallis, Dunn’s multiple comparisons.

Since both Ca^2^+ and Na+ channels have critical roles in controlling the excitability of sensory neurons, we further tested whether NGF increased the activity of voltage-gated ion channels by making patch-clamp recordings under the voltage-clamp mode in dissociated mouse DRG. NGF (50 nM) increased total Ca^2^+ currents (Figure 4E) and current density (Figure 4F, G) by ∼50% when compared to vehicle (DMSO)-treated cells. EG00229 prevented NGF-induced activation of Ca^2^+currents, but had no effect on Ca^2^+currents in unstimulated DRGs (Figure 4F, G). Half-maximal activation and inactivation *(V112)* potential, as well as slope factor values *(k)* for activation and inactivation, were similar between the conditions tested, except for an ∼8-mV hyperpolarizing shift in Ca^2^+channel activation induced by EG00229 when compared to DMSO- and NGF-treated DRGs (Figure 4H, Supplemental Table S2). Similarly, EG00229 normalized NGF-induced increases in Na+ currents (Figure 4l). NGF caused a two-fold increase in Na+ current density (Figure 4J) and peak current density (Figure 4K) when compared to vehicle (DMSO)-treated DRG neurons. EG00229 abolished these effects of NGF but had no effects on Na+ currents in unstimulated DRGs (Figure 4J, K). There were no detectable changes in the voltage-dependence of activation and inactivation between the conditions tested, except for a -10-mV depolarizing shift in the *V112of* inactivation of EG00229 and NGF­ treated cells compared to control (Figure 4L). These data implicate voltage-gated Ca^2^+ and Na+ channels as downstream effectors of NRP1-mediated NGF signaling.

To assess human translation, the contribution of NRP1 to NGF-stimulated activation of dispersed human DRG neurons was similarly studied. Human NGF (50 nM) increased the number of action potentials elicited by a depolarizing ramp pulse (Figure 4M, N). Pre-incubation with the NRP1 inhibitor EG00229 (30 µM) (25) prevented NGF-induced hyperexcitability (Figure 4M, N). NGF or EG00229 did not affect the resting membrane potential (Figure 4O) or the rheobase (Figure 4P).

Thus, NRP1 facilitates NGF-induced sensitization of nociceptors in rodents to humans.

### NRP1 inhibitors suppress NGF-induced nociception in mice

NGF injection is known to induce pain in humans and rodents (5). In addition to sensitizing ion channels, NGF/TrkA signaling promotes chronic pain through the upregulation of genes encoding neuropeptides, ion channels and receptors that mediate pain (e.g., substance P, TRPV1) (5). To test whether NRP1 is necessary for NGF-induced nociception, mouse NGF (50 ng, 10 µl) and NRP1 inhibitors or control reagents were administered to the hindpaws of male mice by intraplantar (i.pl.) co-injection (Figure 5A). Withdrawal responses of injected (ipsilateral) hindpaws to stimulation with van Frey filaments (VFF) and radiant heat were measured to evaluate mechanical allodynia and thermal hyperalgesia, respectively. In control experiments, NGF decreased the withdrawal threshold to VFF stimulation and reduced the latency of withdrawal to thermal stimulation within 30 min for at least 4 h, indicating mechanical allodynia and thermal hyperalgesia (Figure 5B-1). EG00229 (1, 10 or 30 µM/10 µl) (25) exhibited dose­ dependent effects on NGF-induced mechanical allodynia and thermal hyperalgesia compared to vehicle control (PBS) (Figure 5B, F, Supplemental Figure S2A, B). EG00229 at 30 µM strongly inhibited mechanical allodynia and thermal hyperalgesia for 2 h, whereas 10 µM EG00229 inhibited only thermal hyperalgesia at 1 h, and 1 µM EG00229 had no effect. CendR (0.2, 2, 10 µM/10 µl) also had dose-dependent effects on NGF-induced mechanical allodynia and thermal hyperalgesia compared to control (Ctrl) peptide (Figure 5C, G, Supplemental Figure S2C, D). CendR at 10 µM strongly inhibited mechanical allodynia and thermal hyperalgesia for 2 h, 2 µM CendR inhibited mechanical allodynia for 4 h and thermal hyperalgesia for 1 h, whereas 0.2 µM CendR had no effect. Compound 5 (Cpd-5; 30 µM/10 µl), which like EG00229 blocks VEGF-A interaction with NRP1 (36), inhibited NGF-induced mechanical allodynia and thermal hyperalgesia for 1 h compared to vehicle control (Figure 5D, H). Vesencumab (7 µg/10 µl) (16) inhibited NGF-induced mechanical allodynia for 2 h and thermal hyperalgesia for 1 h compared to lgG control (Figure 5E, I). Measurement of the integrated withdrawal responses (area under curve (AUC) of time courses) confirmed the inhibitory actions of these four mechanistically distinct NRP1 inhibitors on NGF-induced nociception (Figure 5J, K). None of the inhibitors affected the withdrawal responses of the ipsilateral hindpaws to mechanical or thermal stimuli in mice that did not receive NGF (Figure 5B-I). NGF injection into the hindpaws of female mice caused a comparable degree of mechanical allodynia and thermal hyperalgesia as in male mice, and EG00229 similarly inhibited NGF-induced nociception in female and male mice (Figure 5L, M).

**Figure 5.**
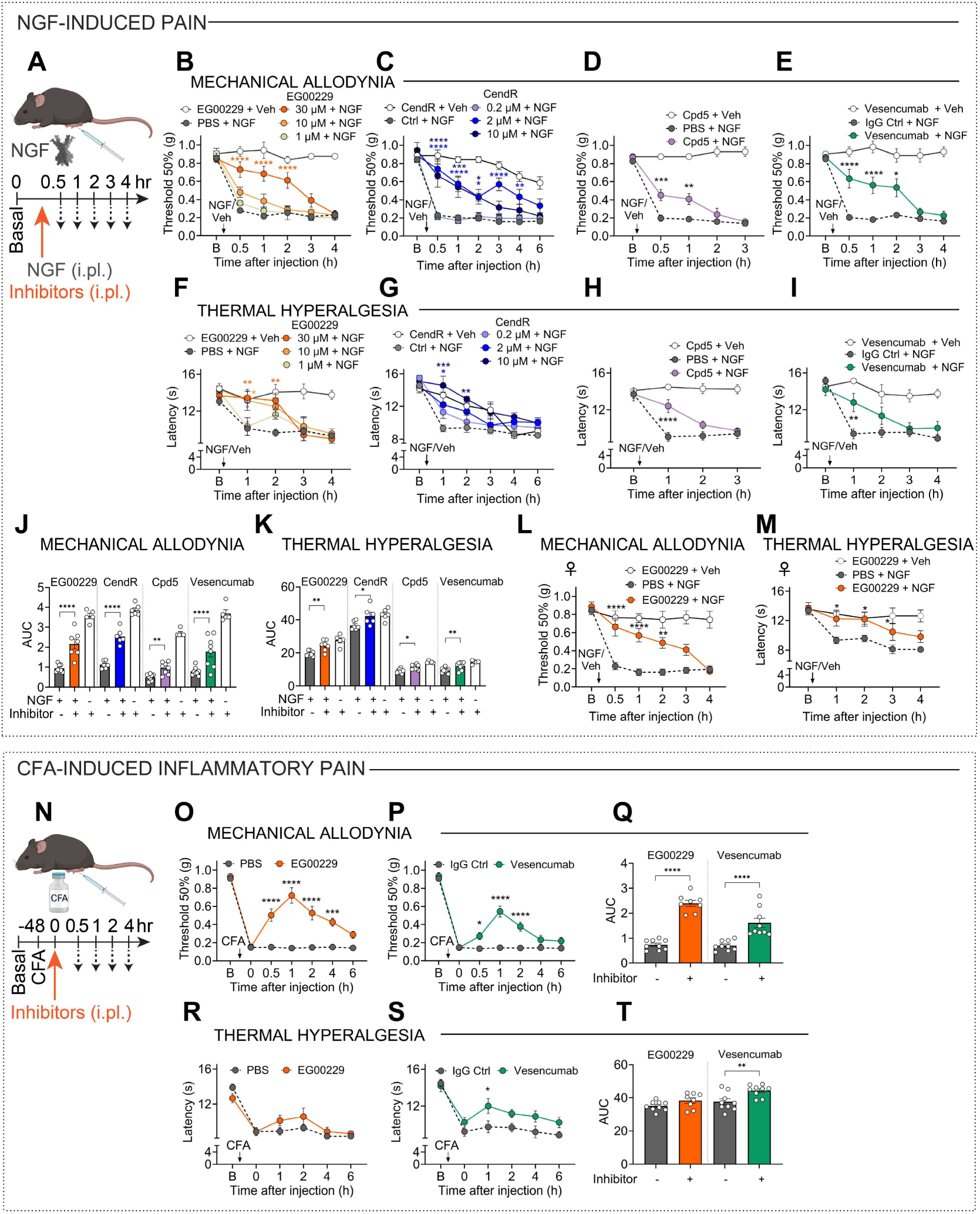
NRP1 inhibition abrogates NGF- and CFA-induced pain in mice. **A-M.** NGF-induced pain in male **(B-K)** and female **(L, M)** mice. Effects of NRP1 inhibitors EG00229 **(B, F;** 1, 10, 30 µM/10 µl i.p1.), CendR **(C, G;** 0.2, 2 and 10 µM/10 µli.pI.) or Compound 5 **(D, H;** Cpd5; 30 µM/10 µli.pI.) or an antibody against the b1 domain of NRP1, Vesencumab **(E, I;** 7 µg/10 µl i.pI.) in male mice. After baseline (’’B”) measurements, inhibitors were co-injected with mouse NGF (50 ng/10 µl, i.p1.). Mechanical allodynia **(B-E)** and thermal hyperalgesia **(F-I)** were measured. N=5-8 male mice per group. **J, K.** Area under curve (AUC) of EG00229 (30 µM/10 µl), CendR (2 µM/10 µl), Compound 5 (30 µM/10 µl) and Vesencumab (7 µg/10 µl) time courses. **L, M.** Effects of NRP1 inhibitor EG00229 (30 µM/10 µl i.pI.) on NGF-induced nociception in female mice. Mechanical allodynia **(L)** and thermal hyperalgesia **(M)** were measured. N=5-8 female mice per group. **N-T.** CFA-induced inflammatory pain. Effects of EG00229 **(0, R,** 30 µM/10 µl) or Vesencumab **(P, S,** 7 µg/10 µl). Inhibitors were injected (i.pl.) 48 h after CFA (i.pl.). Q, T. Area under curve of time courses. Mechanical allodynia **(O-Q)** and thermal hyperalgesia **(R-T)** were measured. N=8-9 mice per group. Mean±SEM. *P<0.05, **P<0.01, ***P<0.001, ****P<0.0001 vs. PBS, control (Ctrl) peptide or lgG Ctrl. B-I, L, M, 0, P, R, S. 2-way ANOVA, Sfdak’s multiple comparisons. J, K, Q, T. 1-way ANOVA, Dunnett multiple comparisons.

NGF mediates inflammatory pain (5), providing an opportunity to determine whether NRP1 is necessary for the pronociceptive actions of endogenous NGF. Complete Freund’s Adjuvant (CFA, 1 mg/ml, 10 µl, i.pl.) was injected into the hindpaws of male mice. After 48 h, EG00229 (30 µM/10 µl) or vehicle (PBS, 10 µl, i.pl.) or Vesencumab mAb (7 µg/10 µl) or lgG Ctrl (7 µg/10 µl) was injected into the inflamed hindpaw (Figure 5N). CFA induced sustained mechanical allodynia and thermal hyperalgesia (Figure 5N-T). EG00299 reversed mechanical allodynia for 4 h (Figure 5O) but did not affect thermal hyperalgesia (Figure 5R). Vesencumab reversed mechanical allodynia for 2 h (Figure 5P) and thermal hyperalgesia for 1 h (Figure 5S). Measurement of the integrated withdrawal responses confirmed these inhibitory effects (Figure 5Q, T).

The finding that four NRP1 inhibitors (EG00229, CendR, Cmpd5, Vesencumab) suppress NGF­ evoked nociception supports the hypothesis that NRP1 enhances the signaling competency of the NGF­ TrkA complex that induces pain.

### NRP1 controls NGF and TrkA kinase signaling

NGF primarily signals through TrkA (29), where NGF binds to extracellular immunoglobulin-like domains to induce conformational changes throughout the RTK dimer that initiate auto- and trans­ phosphorylation of intracellular tyrosine residues (e.g., Y490, Y785). To determine whether NRP1 controls NGF-stimulated phosphorylation of TrkA, a proximal and necessary component of TrkA signaling, dissociated mouse DRG neurons were stained with a phospho-specific antibody to TrkA phosphorylation at residue Y785 (equivalent to Y791 in human TrkA). NGF (100 nM, 15 min) stimulated a -1 .4-fold increase in the intensity TrkA phospho-Y785 immunofluorescence compared to vehicle (Figure 6A, B). Pre­ incubation with EG00229 (30 µM, 30 min) prevented NGF-stimulated phosphorylation of TrkA but had no effect on the basal unstimulated phosphorylation. These results support the hypothesis that NRP1 is necessary for NGF-induced activation of TrkA.

**Figure 6.**
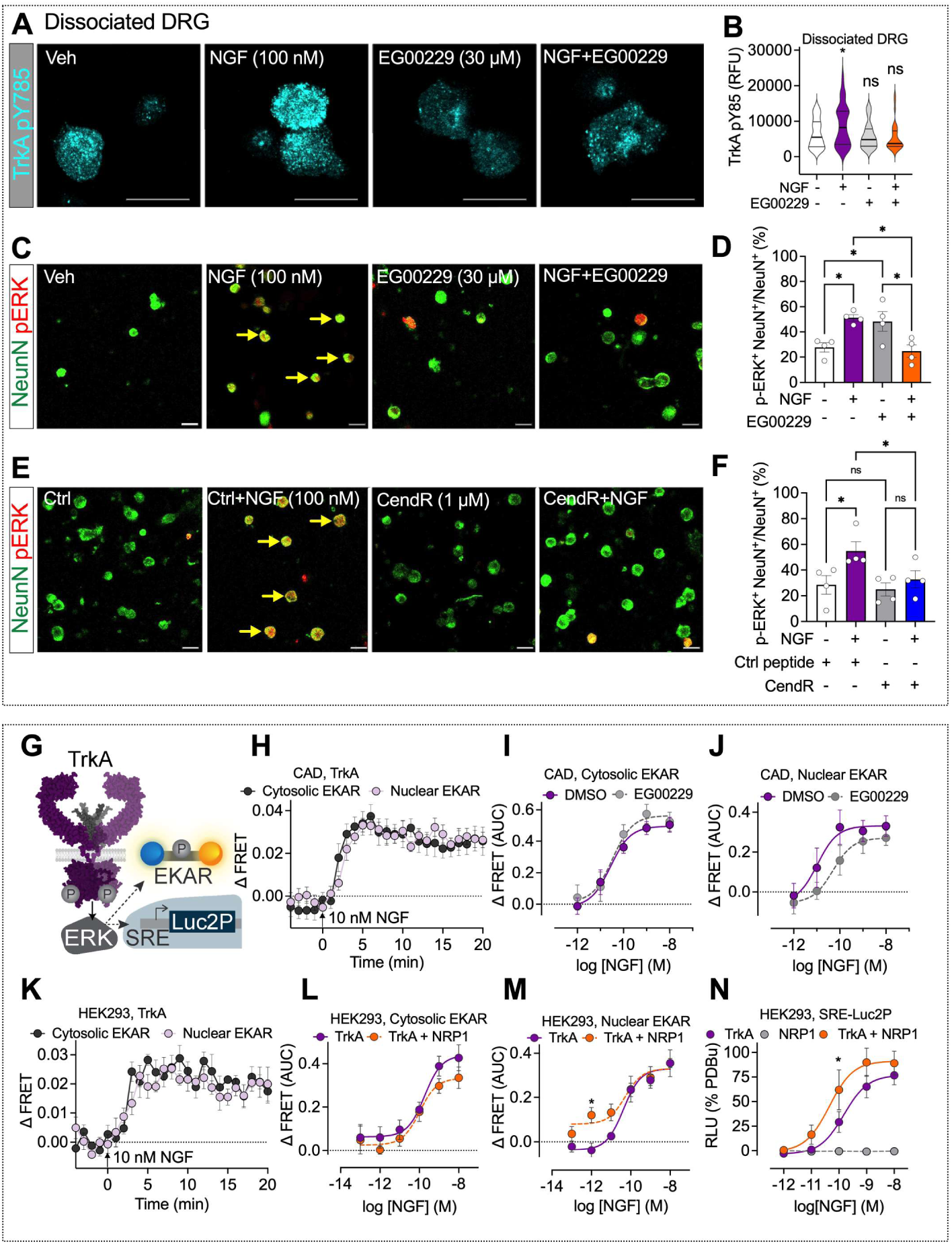
NRP1 modulates TrkA-mediated kinase signaling. **A-B.** Effect of mouse NGF (100 nM, 15 min) and NRP1 inhibitor EG00229 (30 µM, 30 min pre-incubation) on phosphorylated TrkA Y785 staining in mouse DRG neurons. N=34-44 neurons from 3 independent experiments. Scale bar, 20 µm. **C-F.** Effect of mouse NGF (100 nM) and NRP1 inhibitors EG00229 (30 µM, 30 min pre-incubation) **(C, D)** and CendR (1 µM, 30 min pre-incubation) **(E-F)** on phosphorylated ERK Thr202/Tyr204 staining in mouse DRG neurons. N=15-432 neurons from 4 independent experiments. Scale bar, 20 µm. **G-N.** NGF-induced ERK signaling measured using FRET-based EKAR biosensors **(H-M)** or a downstream luciferase reporter **(N). H-M.** NGF-induced modulation of ERK activity using biosensors localized to the cytosol **(H,** I) or nucleus **(H, J)** in neuron-like CAD cells expressing human TrkA. Kinetics of NGF-induced ERK monitored in CAD cells **(H),** comparing increasing NGF concentrations after pre-incubation with EG00229 (30 µM, 30 min) **(I­ J). K-M.** ERK signaling in HEK293T cells expressing TrkA alone or expressing both TrkA and NRP1 **(L, M). N.** Effect of increasing NGF concentrations on ERK transcription in cells expressing TrkA, NRP1 or both (% positive control, 10 µM PDBu). RFU, relative fluorescence units. AUC, area under curve. Data from 4-8 independent experiments with triplicate wells. Mean±SEM. *P<0.05, ***P<0.001, ****P<0.0001. B, M, N. 1-way ANOVA, Sfdak’s multiple comparisons. D, F. 2-way ANOVA, Tukey’s multiple comparison.

Activated TrkA stimulates extracellular signal-regulated kinase (ERK) phosphorylation, as well as AKT and phospholipase Cy. In addition to triggering neuronal development, TrkA-induced ERK signaling contributes to NGF-induced sensitization of nociceptors and pain (5). To determine whether NRP1 is necessary for NGF-induced activation of ERK, dissociated mouse DRG neurons were co-incubated with NGF (100 nM, 30 min) or vehicle together with inhibitors of NRP1 or control reagents. Cultures were stained with antibodies to phosphorylated T202/Y204 ERK1/2 and to the neuronal marker NeuN. NGF stimulated a ∼1.8-fold increase in the number of neurons expressing phosphorylated ERK1/2 (Figure 6C-F, yellow arrows). EG00229 (30 µM, 30 min) or CendR (1 µM, 30 min) prevented NGF-stimulated phosphorylation of ERK1/2 in neurons (Figure 6D, F). Thus, NRP1 is necessary for NGF stimulation of ERK1/2 activity in DRG neurons.

The contribution of NRP1 to ERK activation was monitored in CAD cells, a neuron-like cell line modified from the Gath.a catecholaminergic cell line obtained from a mouse tumor (37). While CAD cells lack TrkA expression, they express p75NTR and NRP1 (Supplemental Figure S3). To probe NGF/TrkA ERK signaling with high spatial and temporal resolution, Forster resonance energy transfer (FRET) EKAR biosensors targeted to the cytosol or nucleus were coexpressed with human TrkA in CAD cells. EKAR biosensors contain a reversible substrate sequence separated by two fluorophores (Figure 6G). NGF activated ERK in the cytosol and nucleus within 5 min for at least 20 min (Figure 6H). NGF-induced ERK activation was concentration-dependent, with a higher potency for nuclear than cytosolic ERK (Figure 6l, J, Supplemental Table S3). Pre-incubation with EG00229 (30 µM) did not affect NGF activation of cytosolic ERK (Figure 6l), but significantly reduced the potency for NGF activation of nuclear ERK (Figure 6J, Supplemental Table S3; NGF ECso: DMSO control, ∼8 pM; EG00229 ∼98 pM; paired t-test, P=0.023).

The contribution of NRP1 to NGF-induced ERK signaling was also investigated by NRP1 overexpression with TrkA in HEK293T cells, which express low endogenous levels of these proteins. As in CAD cells, NGF activated ERK in the cytosol and nucleus of HEK293T cells (Figure 6K). While NRP1 coexpression did not influence cytosolic ERK signaling (Figure 6L), NRP1 expression significantly enhanced activation of nuclear ERK in response to low (1 pM) NGF concentrations (Figure 6M). The outcomes of nuclear ERK signaling were further studied by expression of a transcriptional luciferase reporter, where a luminescent protein is produced downstream of an ERK promoter (Figure 6G). NGF stimulated concentration-dependent ERK transcriptional activity with significantly higher potency in cells overexpressing TrkA and NRP1 compared to cells expressing TrkA alone (Figure 6N, Supplemental Table S3; NGF ECso: TrkA alone, 214 pM; TrkA + NRP1, 71 pM; paired t-test, P=0.0004). NGF did not stimulate ERK transcriptional activity in cells expressing NRP1 alone, confirming lack of inherent signaling capability of NRP1 (Figure 6N). These results support the notion that NRP1 enhances NGF-induced TrkA activation and signaling by pathways that underpin nociception. We next sought to determine the mechanism of NRP1-mediated potentiation of NGF/TrkA signaling.

### NRP1 is a chaperone that forms a complex with TrkA to control trafficking from the biosynthetic pathway to the plasma membrane

In addition to directly binding NGF, NRP1 could amplify NGF/TrkA signaling by receptor/co-receptor interactions with TrkA, akin to its co-receptor function with VEGFR2 (38). Cell surface expression of human TrkA and NRP1 was measured by specific substrate-based labeling of enzymatic tags genetically fused to the extracellular N-terminus of either receptor. SnapTag-TrkA and HaloTag-NRP1 were expressed in HEK293T and CAD cells and covalently labeled with membrane-impermeant AlexaFluor® substrates, thereby selectively labeling receptors transported to the cell surface. SnapTag-TrkA and HaloTag-NRP1 were highly co-localized at the cell surface of HEK293T and CAD cells, with the latter also showing a high level of co-localization in subcellular compartments (Figure 7A). Labeling specificity was confirmed in cells expressing TrkA or NRP1 alone (Supplemental Figure S4).

**Figure 7.**
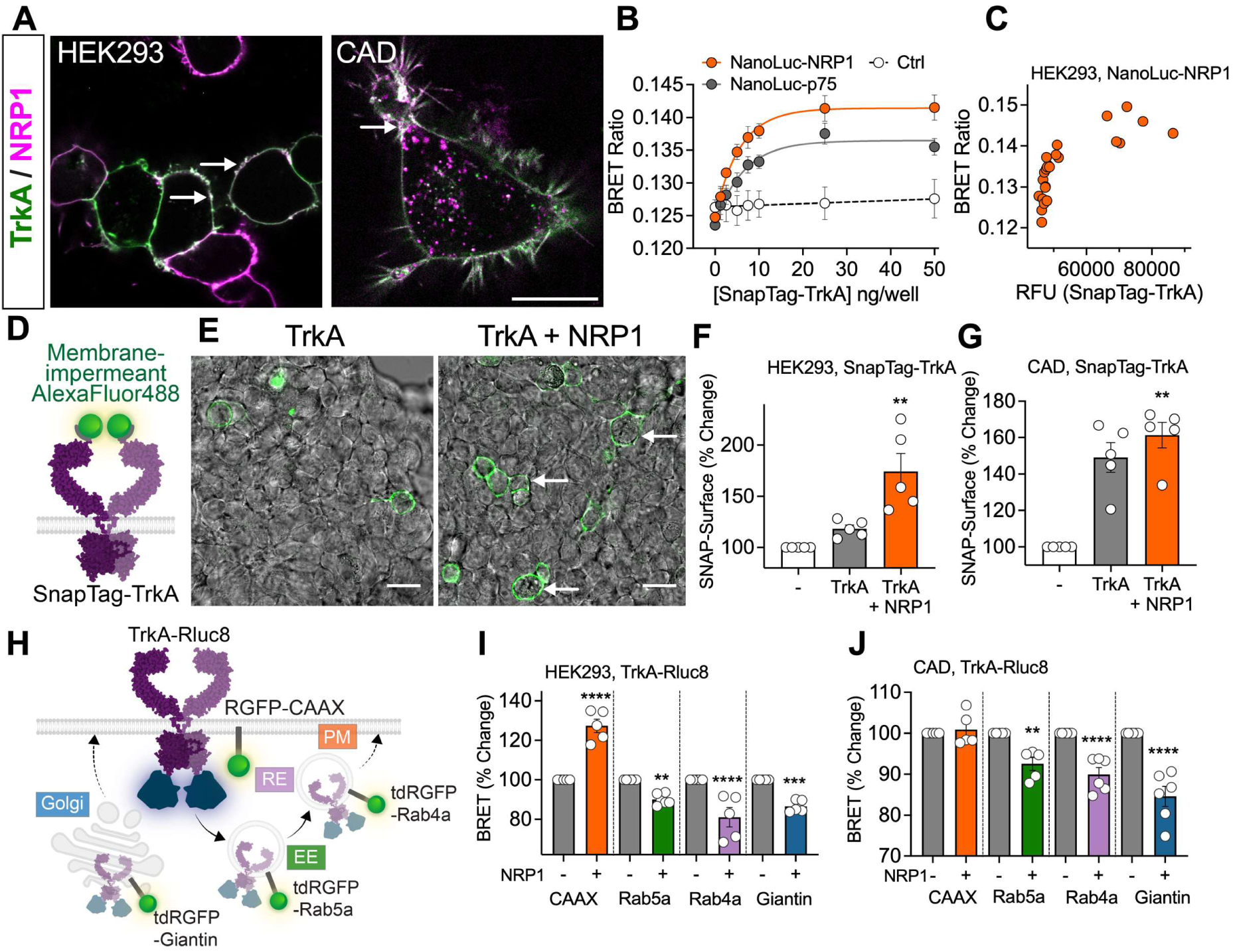
TrkA and NRP1 form a heteromeric complex. **A.** HEK293T or CAD cells expressing SnapTag­ TrkA and HaloTag-NRP1 simultaneously labeled with membrane-impermeant substrate (SNAPTag-Alexa Fluor® 488, HaloTag-Alexa Fluor® 660). Representative images from N=S independent experiments. **B, C.** BRET assays to monitor proximity between Nanoluc-NRP1 or Nanoluc-p75^NTR^ and increasing SnapTag­ TrkA DNA. Negative control, Nanoluc-TrkA and SnapTag-CALCRL. Representative replicate **(C)** plotting BRET against relative fluorescence units (RFU). **D-G.** Cell-surface TrkA in HEK293T imaged in the absence or presence of NRP1 **(E). F-G.** Quantified fluorescence without receptor (-), SnapTag-TrkA alone or SnapTag-TrkA co-transfected with NRP1 in HEK293T **(F)** or CAD **(G)** cells. **H-J.** BRET between TrkA tagged with *Renilla* luciferase (Rluc8) and *Renilla* green fluorescent protein (RGFP) tagged markers of the plasma membrane (PM, RGFP-CAAX), early endosome (EE, tdRGFP-RabSa), recycling endosomes (RE, tdRGFP-Rab4a) or the cis-Golgi apparatus (tdRGFP-Giantin). HEK293T cells (I) or CAD cells **(J)** were transfected with TrkA-Rluc8 in the absence(-) or presence(+) of NRP1. BRET was normalized relative to TrkA-Rluc8 alone (100%). Scale bar, 20 µm. Data from 5-6 independent experiments with triplicate wells. Mean±SEM. *P<0.05, **P<0.01, ***P<0.001, ****P<0.0001. F, G. Paired t-test. I, J. 1-way ANOVA with Sfdak’s multiple comparisons.

To investigate the formation of a heteromeric complex between TrkA and NRP1, BRET was measured in HEK293T cells expressing Nanoluc-NRP1 and SnapTag-TrkA. Extracellular N-terminal Nanoluc acts as an energy donor to excite a nearby SnapTag. Measurement of BRET between Nanoluc­ NRP1 and increasing levels of SnapTag-TrkA revealed a hyperbolic relationship, indicative of assembly of a heteromeric complex between Nanoluc-NRP1 and SnapTag-TrkA (Figure 7B, C). As a positive control, a hyperbolic BRET signal relationship was also detected between Nanoluc-p75NTR, which is known to interact with TrkA (5), and increasing levels of SnapTag-TrkA. In contrast, there was a linear BRET signal between Nanoluc-NRP1 and increasing levels of SnapTag-calcitonin-like receptor (CALCRL), an unrelated transmembrane receptor for CGRP. To verify whether this relationship was observed with protein expression, data from a representative experiment quantifying cell surface SnapTag labeling also demonstrated a hyperbolic curve between TrkA and NRP1 (Figure ?C). These results suggest that NRP1 and TrkA co-localize at the plasma membrane as a heteromeric complex.

Recruiting TrkA to the plasma membrane would enable increased access to extracellular NGF. The effect of NRP1 on the cell surface expression of TrkA was determined using the membrane-impermeant SnapTag fluorophore to selectively label cell surface TrkA (Figure 7D). NRP1 coexpression increased level of TrkA at the surface in HEK293T cells by 150 ± 43% and of CAD cells by 109 ± 4% (Figure 7E-G). Enhanced bystander BRET, which capitalizes on the endogenous affinity of pairs of *Reni//a-tagged* proteins to boost sensitivity (39), was used to quantify the effects of NRP1 expression on the localization of TrkA in different subcellular compartments of living cells. TrkA tagged on the C-terminus with Renilla luciferase (Rluc8, BRET) was coexpressed in HEK293T or CAD cells with proteins resident of the plasma membrane (CAAX), early endosomes (Rab5a), recycling endosomes (Rab4a) and the cis-Golgi apparatus (Giantin) tagged with *Renilla* green fluorescent protein (RGFP) (Figure 7H). In HEK293T cells, NRP1 expression significantly increased BRET between TrkA-Rluc8 and RGFP-CAAX and significantly decreased BRET between TrkA-Rluc8 and tdRGFP-Rab5a, tdRGFP-Rab4a and tdRGFP-Giantin (Figure 7l). In CAD cells, NRP1 expression did not affect BRET between TrkA-Rluc8 and RGFP-CAAX but significantly decreased BRET between TrkA-Rluc8 and tdRGFP-Rab5a, tdRGFP-Rab4a and tdRGFP-Giantin (Figure 7J). These results indicate that NRP1 expression causes a redistribution of TrkA from subcellular regions involved in receptor recycling or *de nova* export to the plasma membrane, consistent with a chaperone function. Control studies confirmed NRP1 was successfully expressed using a HaloTag label (Supplemental Figure S5A-B). While there was a small reduction TrkA-Rluc8 expression in CAD cells upon NRP1 coexpression, NRP1 coexpression had no significant effect on overall TrkA expression in HEK293T cells (Supplemental Figure S5C-D). These results suggest that NRP1 acts as a chaperone for TrkA.

### NRP1 enhances NGF-induced TrkA trafficking and dimerization

Upon NGF stimulation, TrkA traffics to endosomes by clathrin-dependent and -independent mechanisms. NGF/TrkA signalosomes are retrogradely transported in endosomes or multivesicular bodies to the soma of sympathetic neurons, where they mediate the neurotrophic actions of NGF (3, 4). To determine the effects of NRP1 on NGF-evoked endocytosis of TrkA, BRET was measured between TrkA­ Rluc8 and plasma membrane marker RGFP-CAAX (Figure 8A). NGF caused a concentration-dependent decrease in TrkA-Rluc8 and RGFP-CAAX BRET (EC50 ∼870 pM), with 10-100 nM NGF inducing maximal decrease within 5 min that was sustained for 20 min, consistent with TrkA endocytosis (Figure 8B, Supplemental Table S3). Hypertonic (0.45 M) sucrose or the clathrin inhibitor pitstop 2 (30 µM) prevented NGF-induced endocytosis of TrkA (Figure 8C, D). Coexpression of NRP1 with TrkA enhanced NGF­ stimulated endocytosis of TrkA (Figure 8E, F). This effect of NRP1 was observed at higher NGF concentrations (>10 nM), with minimal effect on the potency of NGF-induced endocytosis of TrkA (Supplemental Table S3). NGF also stimulated concentration-dependent removal of TrkA from the plasma membrane of CAD cells (EC50 ∼490 nM; pEC50 = 9.31 ± 0.18, N=4) that was inhibited by hypertonic sucrose and pitstop 2 (Figure 8G-J). Knockdown of endogenous NRP1 with siRNA significantly inhibited NGF­ induced endocytosis of TrkA in CAD cells (Figure 8K, L). NRP1 knockdown was confirmed at a protein level using immunofluorescence in CAD cells, while there was no effect on TrkA expression (Supplemental Figure S5E-F). Thus, whereas NRP1 overexpression enhances agonist-stimulated endocytosis of TrkA in HEK293T cells, NRP1 knockdown has the opposite effect in CAD cells. The results are consistent with a chaperone role for NRP1 in NGF-stimulated endocytosis of TrkA.

**Figure 8.**
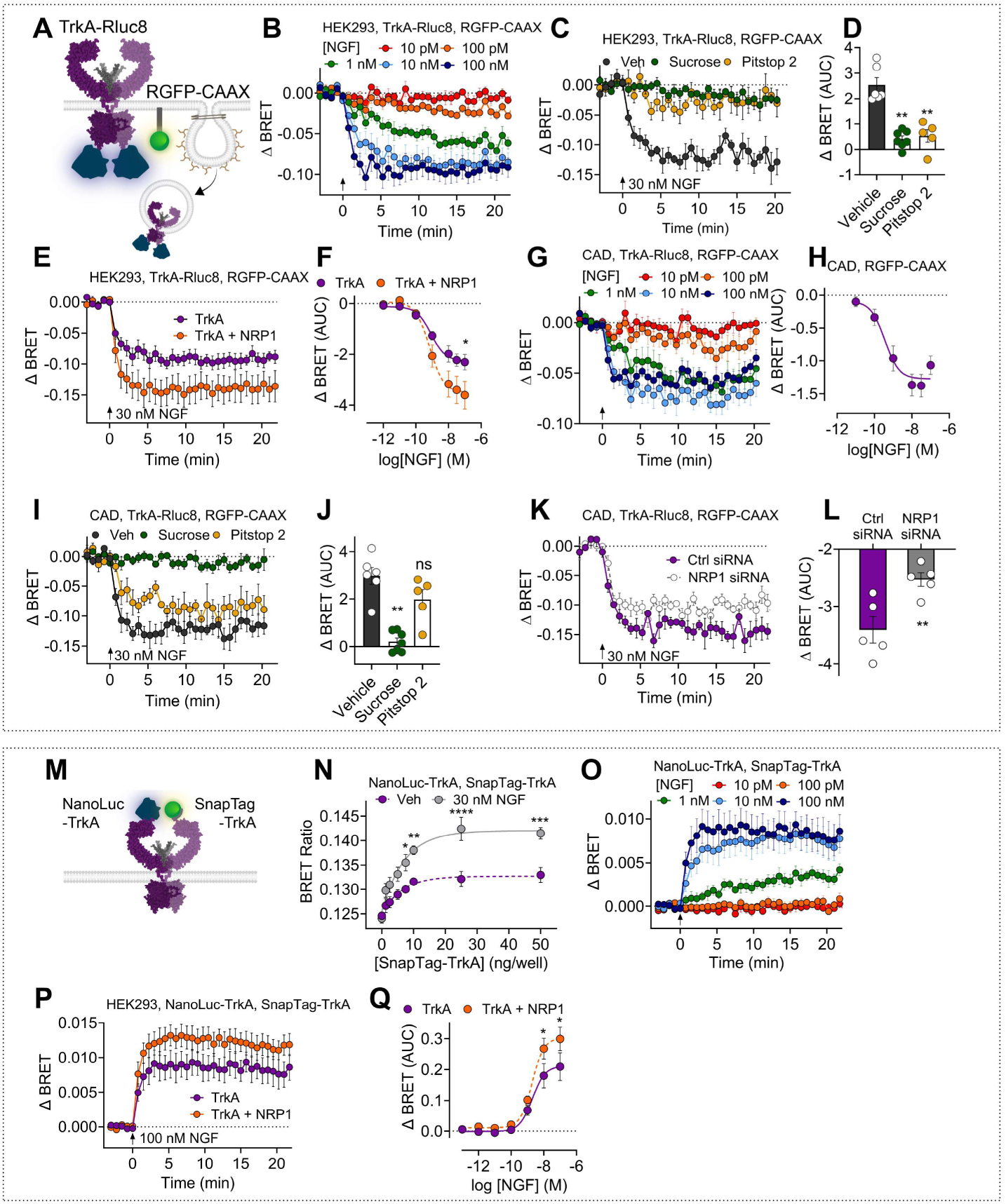
NRP1 modulates NGF-induced TrkA trafficking and RTK oligomerization. **A-L.** BRET measurements between Rluc8-tagged TrkA and a fluorescent marker of the plasma membrane (RGFP­ CAAX) at 37°C. Decreased BRET indicates TrkA-Rluc8 removal from the plasma membrane **(A). B-F.** Trafficking in HEK293T cells incubated with graded concentrations of NGF **(B),** or pre-incubated for 30 min with hypertonic sucrose (0.45 M), clathrin inhibitor pitstop 2 (30 µM) or vehicle (Veh) **(C, D)** before NGF stimulation. Effect of NRP1 coexpression on NGF-stimulated endocytosis of TrkA **(E, F). G-L.** Trafficking in CAD cells expressing human TrkA incubated with graded concentrations of NGF **(G, H),** or pre-incubated for 30 min with hypertonic sucrose (0.45 M), clathrin inhibitor pitstop 2 (30 µM) or vehicle (Veh) **(I, J)** before NGF stimulation, or transfected with control or mouse NRP1 siRNA **(K, L). M-Q.** BRET measurements between Nanoluc-TrkA and SnapTag-TrkA to assess TrkA oligomerization in HEK293T cells. **N.** BRET with increasing expression of SnapTag-TrkA and fixed Nanoluc-TrkA (10 ng), after 30 min incubation with vehicle or NGF. **O-Q.** Using a fixed donor:acceptor ratio (1:2.5), oligomerization kinetics at 37°C in response to increasing NGF concentrations (**O**) in the absence and presence of NRP1 coexpression **(P, Q;** showing the same 100 nM NGF kinetic data for TrkA in **O** and **P).** AUC, area under curve. Data from 4-6 independent experiments with triplicate wells. Mean±SEM. *P<0.05, **P<0.01, ***P<0.001, ****P<0.0001. G, Unpaired t-test. D, I, K. 1-way ANOVA with Sidak’s multiple comparisons.

Dimerization of TrkA is pivotal for NGF/TrkA signaling (5). The contribution of NRP1 to TrkA oligomerization was evaluated by measuring BRET between Nanoluc-TrkA and SnapTag-TrkA in HEK293T cells (Figure BM). Expression of increasing amounts of SnapTag-TrkA in HEK293T cells expressing a fixed amount of Nanoluc-TrkA produced a hyperbolic BRET signal, indicating oligomerization (Figure 8N). NGF (30 nM) significantly enhanced this signal, consistent with agonist-evoked TrkA oligomerization. In cells expressing a fixed ratio of SnapTag-TrkA and Nanoluc-TrkA, NGF stimulated a concentration-dependent increase in BRET that was maximal after 5 min and sustained for at least 20 min (ECso 2.3 nM; Figure 8O, Supplemental Table S3). While NGF had a slightly lower potency with respect to dimerization than endocytosis, the kinetics of TrkA dimerization and endocytosis were similar. NRP1 overexpression enhanced TrkA oligomerization in response to higher NGF concentrations (>10 nM) (Figure BP, Q). As dimerization is the first step in RTK activation, with well-established evidence for TrkA signaling from endosomes, these results provide a mechanistic insight into the role of NRP1 in controlling TrkA signaling and trafficking.

### GIPC1 mediates NRP1/TrkA interactions and NGF-induced pain signaling

While NRP1 lacks intrinsic catalytic activity, the short cytoplasmic C-terminus interacts with G Alpha Interacting Protein C-terminus 1 (GIPC1), or synectin, through a PDZ domain (40). GIPC1 also interacts with the membrane-proximal regions of TrkA (41). Thus, interaction with GIPC1 could underpin the co­ receptor function of NRP1 in NGF/TrkA-evoked nociception. Analysis by RNAScope^TM^ *in situ* hybridization revealed that *Gipc1* mRNA was expressed by all mouse DRG neurons (Figure 9A). In humans, *Gipc1* was expressed by 100% of Ntrk1-positive neurons and *Ntrk1* was expressed by 85% of Gipc1-positive neurons (Figure 9B, C).

**Figure 9.**
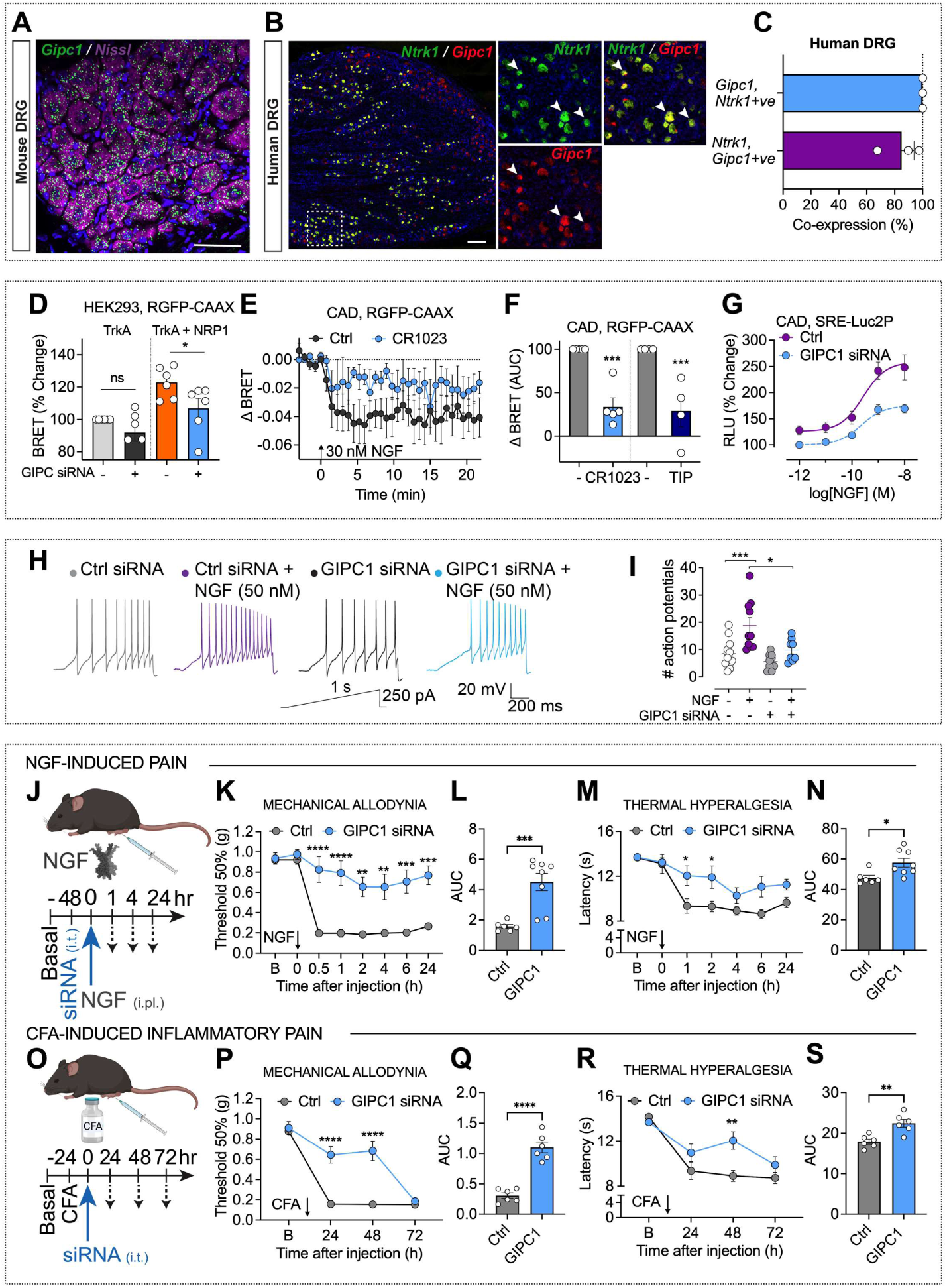
GIPC1 modulates TrkA trafficking, signaling and NGF-induced nociception. **A-C.** RNAScope™ localization of *Gipc1* mRNA in mouse DRG **(A)** and of *Ntrk1* and *Gipc1* mRNA in human DRG **(B).** Arrows indicate mRNA expression within the same cell. Representative images, N=5 mice and N=3 humans. Scale bar, 500 µm. C. Percentage of human DRG neurons expressing *Ntrk1 or Gipc1* that coexpress *Gipc1* or *Nrp1.* Hybridized positive neurons(%) from N=3 humans. **D.** Effect of GIPC1 siRNA on BRET measurements of TrkA levels at the plasma membrane of HEK293T cells under basal conditions and after coexpression with NRP1. **E, F.** Effect of 30 min pre-incubation of GIPC1 antagonist (300 µM CR1023 or inactive control, Ctrl) or myosin VI inhibitor (50 µM 2,4,6-Triiodophenol, TIP) on NGF-induced TrkA-Rluc8 trafficking from a marker of the plasma membrane (RGFP-CAAX) in CAD cells. **G.** Effect of GIPC1 siRNA on NGF-induced downstream ERK transcription in CAD cells. Data from 5-6 independent experiments with triplicate wells. **H,** I. Sample traces of action potential firing in mouse DRG neurons evoked by injecting a 1 s ramp pulse from O to 250 pA **(G),** with the number of evoked action potentials **(H).** N=?-10 cells. **J-N.** NGF-induced pain. Effects of GIPC1 or Ctrl siRNA (i.t.) on NGF (50 ng/10 µl, i.pl.)-induced mechanical allodynia (K, L) and thermal hyperalgesia **(M, N)** in the ipsilateral paw. **L, N.** Area under curve of time courses. **0-S.** CFA-induced pain. Effects of GIPC1 or Ctrl siRNA (i.t.) on CFA (i.pl.)-induced mechanical allodynia **(P, Q)** and thermal hyperalgesia **(R, S). Q, S.** Area under curve (AUC) of time courses. N=6-8 mice per group. RLU, relative luminescence units. AUC, area under curve. ‘B’, basal. Mean±SEM. P<0.05, **P<0.01, ***P<0.001, ****P<0.0001. D, F. 1-way ANOVA, Sfdak’s multiple comparisons. I. Tukey’s multiple comparison. K, M, P, R. 2-way ANOVA, Sfdak’s multiple comparisons. L, N, Q, S. Unpaired t-test.

GIPC1 is an intracellular adaptor protein that associates with receptors and channels to regulate their trafficking by interacting with the inwardly-directed myosin VI motor (42). To determine whether GIPC1 is necessary for the TrkA chaperone function of NRP1, BRET between TrkA-Rluc8 and RGFP-CAAX was measured in HEK293T cells after GIPC1 knockdown. GIPC1 siRNA inhibited NRP1-induced plasma membrane expression of TrkA (Figure 9D). In CAD cells, preincubation with a GIPC1 inhibitor (300 µM CR1023 (43)) or a myosin VI inhibitor (50 µM TIP (44)) reduced NGF-induced TrkA trafficking to the plasma membrane (Figure 9E, F). Similarly, GIPC1 siRNA inhibited the maximal response of NGF-induced ERK signaling quantified using the downstream transcriptional reporter in CAD cells, with no effect on potency (Figure 9G) (Ctrl siRNA pECso = 9.58 ± 0.03, GIPC1 siRNA pECso = 9.63 ± 0.09, N=5). GIPC1 siRNA also prevented NGF-induced action potential firing in mouse DRG nociceptors, determined by patch-clamp recordings (Figure 9H, I). GIPC1 siRNA knockdown was confirmed at an mRNA level in HEK293T cells, CAD cells and DRG (Supplemental Figure S6A-C).

To evaluate the role of GIPC1 in NGF-induced nociceptive behavior, GIPC1 or control siRNA was administered to male mice by intrathecal (i.t.) injection 48 h before NGF (50 ng/10 µl, i.pl.) (Figure 9J). In mice treated with control siRNA, NGF caused mechanical allodynia and thermal hyperalgesia in the ipsilateral paw within 30 min for at least 24 h (Figure 9K-N). GIPC1 siRNA prevented NGF-evoked mechanical allodynia for at least 24 h and inhibited thermal hyperalgesia for 2 h. NGF or siRNA administration did not affect withdrawal responses of the contralateral paw to mechanical stimuli (Supplemental Figure S7A). The nociceptive role of GIPC1 was also investigated in a preclinical model of inflammatory pain in male mice. Control or GIPC1 siRNA was administered (i.t.) 24 h after CFA (1 mg/ml, 10 µl, i.pl.) (Figure 9O). In mice treated with control siRNA, CFA caused mechanical allodynia and thermal hyperalgesia for at least 72 h (Figure 9P-S). GIPC1 siRNA inhibited CFA-induced mechanical allodynia and thermal hyperalgesia after 24 and 48 h. CFA or siRNA administration did not affect withdrawal responses of the contralateral paw to mechanical stimuli (Supplemental Figure S7B). GIPC1 siRNA caused knockdown of *Gipc1* mRNA in DRG, determined by RNAScope™ *in situ* hybridization (Supplemental Figure S5D). These results reveal a new role for GIPC1 in NGF-evoked pain.

## Discussion

We report that NRP1 is a previously unrecognized co-receptor for NGF and TrkA signaling of pain, demonstrated in rodent and human tissue. NRP1 inhibition was found to attenuate NGF and TrkA signaling in cell lines and to block the pronociceptive actions of NGF in mice and isolated human and mouse nociceptors, whereas NRP1 overexpression amplified NGF and TrkA signaling and trafficking. In a similar manner, NRP1 is a co-receptor for VEGF-mediated angiogenesis, demonstrated using pharmacological (24) and genetic (45) interventions. Our results show that NRP1 promotes NGF/TrkA-mediated pain by at least two mechanisms: as a co-receptor that interacts with NGF and TrkA to form a ternary NGF/TrkA/NRP1 signaling complex, and as a chaperone that enhances TrkA trafficking from the biosynthetic pathway to the plasma membrane and then to signaling endosomes (Figure 10).

**Figure 10.**
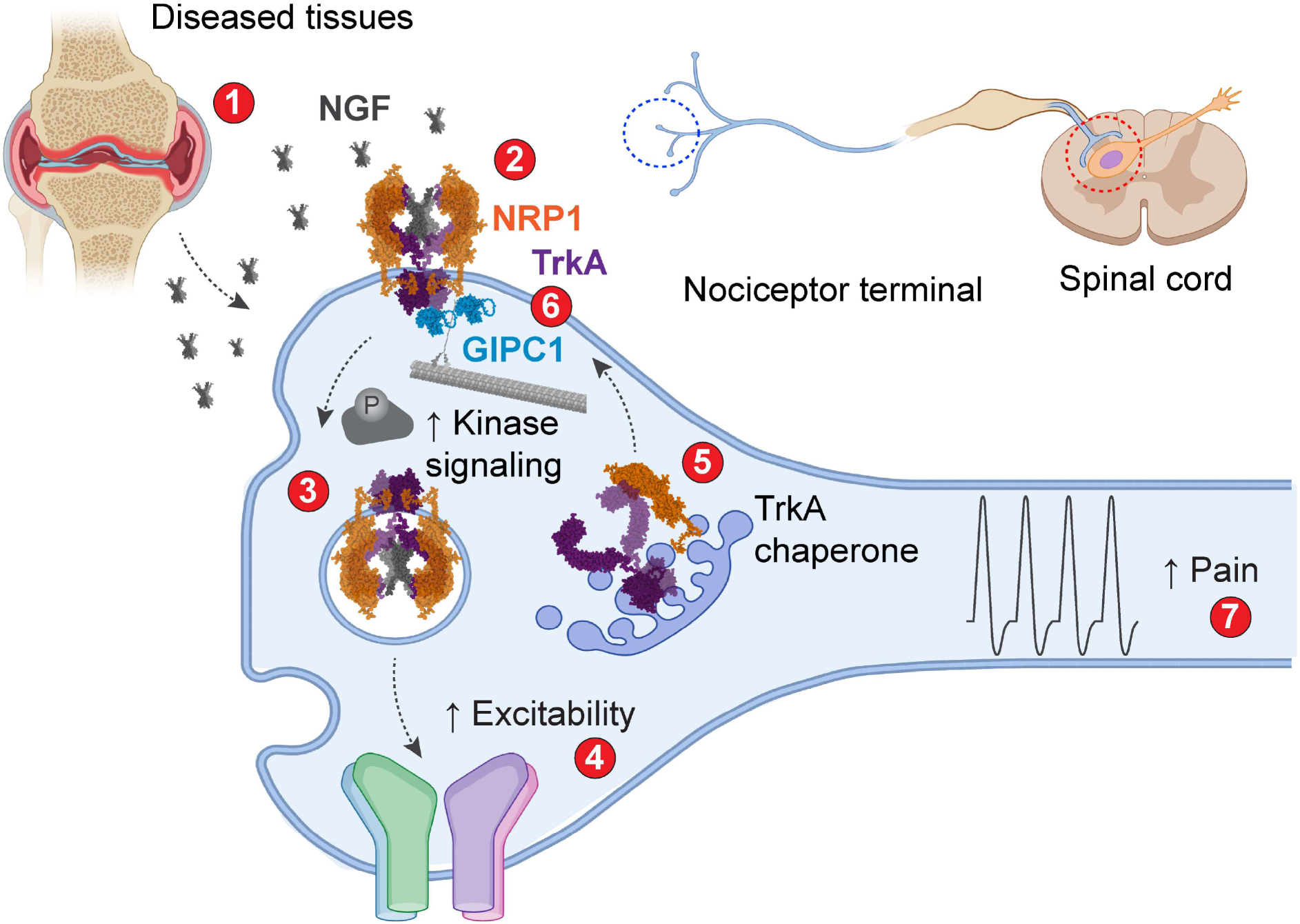
Hypothesized mechanism by which NRP1 mediates NGF/TrkA pain signaling. **1.** NGF is released from diseased tissues (e.g., sites of injury, inflammation, cancer) in close proximity to the peripheral endings of nociceptors. **2.** At the surface of nociceptors, NGF binds to both NRP1 and TrkA, forming a ternary NGF/NRP1/TrkA complex with a 2:2:2 stoichiometry. **3.** TrkA signals from the plasma membrane and endosomes to activate kinases and ion channels. **4.** Activation and sensitization of TRPV1 and Na+ and Ca^2^+channels leads to increased excitability of nociceptors. **5.** NRP1 chaperones TrkA from the biosynthetic pathway to the plasma membrane and to signaling endosomes, which further enhances excitability of nociceptors. **6.** GIPC1 interacts with NRP1 and TrkA, linking the complex to the myosin VI molecular motor to amplify pain signaling. **7.** By binding NGF and interacting with TrkA, NRP1 is a co­ receptor that facilitates NGF/TrkA signaling of pain.

Several observations suggest that NRP1 is an NGF and TrkA co-receptor. *Nrp1/NRP1* was coexpressed with *Ntrk1/TrkA* in peptidergic nociceptors of mouse and human DRG. In support of these findings, a single cell transcriptomics study reported coexpression of *Nrp1* and *Ntrk1* in peptidergic C-fibers in mouse and human DRG, including 37% of CGRP-positive neurons (46). Thus, NRP1 is appropriately colocalized with TrkA in mouse and human nociceptors to control NGF and TrkA signaling of pain. In contrast to our findings, *Nrp1* and *Ntrk1* were not detected in satellite glial cells (46). This discrepancy, which may be attributable to different methodology, warrants further investigation. Although our results provide evidence for NRP1 and TrkA interactions within the same cell, the extracellular domain of NRP1 could interact with the extracellular domain ofTrkA on adjacent cells. In a similar manner, NRP1 can interact with VEGFR2 both when they are coexpressed in the same cell, where the extracellular and intracellular domains associate, and in adjacent cells, where only the extracellular domains associate (47). This intercellular signaling may underlie functional interactions between nociceptors and adjacent cells that contribute to pain (e.g., glial cells).

MST measurements revealed that NGF and NRP1 interact with nanomolar affinity. BRET proximity assays provide evidence that NGF interacts with NRP1 at the cell surface. Future studies of the effects of NRP1 on NGF/TrkA binding kinetics are warranted because NRP1 potentiation of VEGFR2 signaling could be linked to growth factor binding kinetics (26). Measurement of BRET between NRP1 and increasing levels of TrkA revealed a hyperbolic relationship, which is indicative of assembly of a NRP1/TrkA complex. A computational docking protocol, which was based on the structure of the NGF/TrkA complex (29) and on structural features underlying interactions between NRP1 with other growth factors (20, 30–32), predicted the formation of a ternary NGF/TrkNNRP1 complex at the cell surface with a 2:2:2 stoichiometry. This analysis predicts that R118 of the NGF CendR motif K115AVR11s binds to a C-terminal arginine binding pocket of the NRP1 b1 domain through hydrogen bonding interactions between NGF R118 and NRP1 conserved residues Y297, 0320 and Y353. The membrane proximal MAM domain of NRP1 is proposed to orient other extracellular NRP1 domains away from the plasma membrane, enabling one NRP1 molecule to interact with one TrkA molecule and the NGF dimer. Structural analysis of the putative NGF/TrkA/NRP1 complex will be required to confirm this prediction.

The proposed chaperone function of NRP1 is supported by BRET assays showing that NRP1 and TrkA form a heteromeric complex and that NRP1 expression re-routes TrkA from the biosynthetic pathway to the plasma membrane and subsequently to signaling endosomes. By chaperoning TrkA to the cell surface and then to signaling endosomes, NRP1 is expected to amplify the intracellular signaling of NGF/TrkA, which is known to regulate gene transcription (4, 48, 49). The observation that NRP1 expression potentiates NGF-stimulated TrkA dimerization, endocytosis and kinase signaling is consistent with this role of NRP1. Further studies are required to experimentally confirm the potential NRP1/TrkA interaction sites predicted by molecular modeling to mediate the chaperone function of NRP1.

Although TrkA is the principal pronociceptive NGF receptor, p75NTR also contributes to NGF­ induced pain (50). Like NRP1, p75NTR accelerates NGF binding and internalization when coexpressed with TrkA (51). Since the NGF/p75NrR binding site (52) does not directly conflict with the proposed NGF/NRP1 binding site, a TrkA/p75NTR/NRP1 complex may also interact with NGF, although there could be steric clashes between NRP1 and p75NTR for the same binding site on NGF. Further studies are required to determine whether NRP1 regulates NGF interaction with p75NTR_ Despite the glycosylated nature of NRP1, indirect association of NGF with matrix components is unlikely to mediate NGF/NRP1 interactions as NGF does not directly interact with the extracellular matrix (53).

Our results show that GIPC1 is a previously unrecognized mediator of pain. In the context of NGF/TrkA-evoked pain, GIPC1 interacts with TrkA (41) and NRP1 (40), and could thus scaffold TrkNNRP1 interactions to facilitate NGF signaling. By coupling TrkA and NRP1 to the myosin VI molecular motor, GIPC1 may mediate trafficking of TrkA and NRP1 to the plasma membrane and signaling endosomes. In support of this possibility, GIPC1 disruption inhibited NRP1-stimulated translocation of TrkA to the plasma membrane and suppressed NGF-evoked endocytosis of TrkA and ERK signaling, in line with reports in other systems (54). GIPC1 prominently colocalized with NRP1 and thus TrkA in human and mouse nociceptors, and is therefore appropriately located to control NGF-evoked pain. However, since GIPC1 regulates the trafficking of many receptors and ion channels, other mechanisms could also mediate the antinociceptive effects of GIPC1 knockdown in mice.

Electrophysiological and calcium imaging studies of human and mouse nociceptors, and analysis of nociceptive behavior in mice, support a role for NRP1 and GIPC1 in NGF-induced pain. The NRP1 inhibitors (EG00229, CendR, compound 5, Vesencumab) and GIPC1 siRNA suppressed NGF-evoked sensitization of mouse and human nociceptors and mechanical allodynia and thermal hyperalgesia in mice. GIPC1 siRNA prevented NGF-induced nociception in mice. EG00229 had similar effects on mouse and human nociceptors, which supports human translation. Although pharmacological inhibitors can lack selectivity and NRP1 deletion from nociceptors would unequivocally define its role in pain, the finding that mechanistically distinct inhibitors have similar effects provides confidence in specificity. EG00229 and compound 5 inhibit binding of VEGF-A to the b1 domain of NRP1 (25, 36). CendR corresponds to an NGF fragment that includes the two R/KXXR/K domains predicted to interact with the extracellular b domains of NRP1 and likely competes with NGF for NRP1 binding. Vesencumab is a human mAb against the b1b2 domain of NRP1 (16). These inhibitors all suppressed NGF-induced nociception.

NGF causes pain by temporally distinct mechanisms, including rapid sensitization of ion channels and sustained transcription of pronociceptive mediators (55). Our results suggest that NRP1 contributes to both the rapid (TrkA dimerization, phosphorylation, channel activation) and sustained (ERK activation, transcription) actions of NGF **(Figure 10).** The finding that Vesencumab and CendR, which are unlikely to penetrate the plasma membrane, block NGF-evoked nociception, suggest that NRP1 binding to NGF and association with TrkA at the plasma membrane mediate the rapid pronociceptive actions of NGF. By chaperoning TrkA to the plasma membrane and enhancing trafficking of the NGF/TrkA complex to signaling endosomes, NRP1 and GIPC1 could contribute to sustained NGF-evoked nociception. NGF/TrkA signaling endosomes have been identified in sympathetic (49) and DRG (48) neurons, and retrograde NGF/TrkA signaling controls gene expression during the development of sympathetic neurons (49). Whether endosomal signaling of NGF/TrkA contributes to pain, as observed with G protein-coupled receptors (56), deserves further attention, including determination of the contribution of NRP1 to NGF-induced expression of pronociceptive transmitters and channels. NRP1 interacts with semaphorins (19) as a co-receptor for plexins, which regulate TrkA retrograde signaling (57). NRP1 localizes to axonal growth cones (58) and NGF-responsive DRG neurons during neurite sprouting (59). Thus, disruption of NRP1 could affect NGF­ induced neuronal growth, with implications for the neuroplasticity associated with chronic pain.

Further studies are required to determine whether NRP1 inhibitors are a safe and effective treatment for pain. NGF is implicated in inflammatory, neuropathic, surgical and cancer pain (5), highlighting the need to study contributions of NRP1 across pain pathologies. While NGF mAbs provided beneficial pain relief to patients with arthritis, rapidly progressing osteoarthritis in some patients precluded FDA approval

(12). The mechanism responsible for osteoarthritis in patients treated with NGF mAbs is not understood, and whether NRP1 differentially regulates the pronociceptive and protective actions of NGF in joints is unknown. The finding that NRP1 is enriched with TrkA in nociceptors supports targeting NRP1 for the treatment of pain. Therapies targeting interactions between NGF, TrkA and NRP1 in nociceptors, such as CendR peptides designed to block NGF and NRP1 association, may obviate the detrimental effects of global NGF sequestration with mAbs. TrkA and NRP1 are also expressed by chondrocytes, where NGF and TrkA have been implicated in articular cartridge homeostasis (60). Targeting Sema3A-NRP1 signaling has been proposed as a therapy for arthritis (61). The effects of NRP1 antagonism on joint health and disease warrants further study. Because NRP1 interacts with pronociceptive growth factors in addition to NGF, including VEGF-A (18), NRP1 inhibitors could suppress several forms of pain. In the context of cancer, NRP1 inhibitors would be expected to suppress NGF-evoked pain and VEGF-A-mediated angiogenesis in tumors, although impaired wound healing could be a liability. Although NRP1 mAbs were well tolerated in human subjects (62), analysis of side effects of NRP1 inhibition **will** be required to advance NRP1-directed therapies to the clinic. Identifying the antinociceptive efficacy of Vesencumab highlights an opportunity to repurpose a biologic developed for cancer for the treatment of chronic pain, facilitating the development of a non-opioid therapeutic targeting NGF signaling through its novel co-receptor identified in this study.

## Materials and Methods

See Supplemental Materials

### Sex as a biological variable

The role of NRP1 in NGF-evoked nociception was studied in male and female mice.

### siRNA

siRNAs were from Dharmacon (Supplemental Table S4).

### Microscale thermophoresis

Fluorescent NGF was mixed with NRP1 for thermophoresis measurements.

### HiBiT-BRET binding assays

SnapTag-NRP1 or SnapTag-NRP1 Y297A were expressed in HEK293T cells and labeled with SNAP-Surface® Alexa Fluor® 488. Separately, IL-6-HiBiT-VEGF165a or IL-6-NGF­ HiBiT were expressed in other HEK293T cells. Secreted growth factor was collected as cell supernatant. Supernatant was incubated with recombinant HaloTag-LgBiT and nanoluciferase substrate furimazine. NRP1-expressing cells were pre-incubated with vehicle or unlabeled VEGF155a, followed by supernatant with furimazine. Luminescence and fluorescence emissions were recorded.

### NGF/TrkA/NRP1 computational modelling

Ternary complexes of human NGF/TrkA/NRP1 were generated through an information-driven computational docking protocol (27) and were compared to existing biochemical and molecular data.

### Mouse tissue

Anesthetized mice were transcardially perfused with 4% paraformaldehyde. DRG (L4-L5) were fixed in 4% paraformaldehyde, cryoprotected, embedded, and sectioned.

### Human tissue

Donor information has been provided (63). DRG (L4-L5) were fixed in 4% paraformaldehyde, cryoprotected, embedded, and sectioned.

### Lmmunofluorescence

Sections were incubated with primary antibodies to TrkA, NRP1, glutamine synthetase, CGRP, and P2X3. Slides were washed, incubated with fluorescent secondary antibodies and imaged by confocal microscopy.

### RNAScope™ *in situ* hybridization

NRP1, TrkA and GIPC1 were localized by RNAScope™ using probes to mouse and human targets.

### DRG culture

Human DRG suspension cells were from AnaBios Corporation and were studied within 96 h. Thoracic and lumbar mouse DRG were enzymatically dissociated. GIPC1 or Ctrl siRNA plus a plasmid encoding the enhanced green fluorescent protein (eGFP) were expressed by Nucleofector® transfection. Neurons were used within 48 h and siRNA transfection verified by eGFP fluorescence.

### Patch-clamp

Action potentials were recorded in whole-cell patch clamp configuration and current-clamp mode. Action potentials were evoked by a ramp pulse from 0-1000 pA in 1 sec for human DRG, or 0-250 pA in 1 sec for mouse DRG. Rheobase was measured by injecting currents from O pA with an increment of 50 pA in 100 ms (human) or 10 pA in 50 ms (mouse). Ca^2^+currents (ICa^2^+) and Na+ (INa+) currents were recorded as described (17, 18). DRG neurons were preincubated with EG00229 or vehicle (0.1% DMSO) and then challenged with NGF.

### Calcium imaging

DRG neurons were incubated with Fluo-4AM (1 µM) to measure changes in [Ca^2^+)i. Neurons were continuously perfused (10 ml/min) with calcium buffer and maintained at 37°C. Fluorescence was recorded in individual neurons. After basal recording for 1 min, neurons were challenged with capsaicin (100 nM) at 1 min and at 7 min. Depolarization was evoked with 50 mM KCI at 13 min. Neurons were exposed to mouse NGF (100 nM) or vehicle (control) at 5 min. In some experiments, neurons were preincubated with EG00229 (3, 10, and 30 µM) or vehicle (0.1% DMSO, control) for 1 min before capsaicin. CendR or control peptide (0.1, 0.3, and 1 µM) and Vesencumab mAb or control lgG (0.7 µg/ml) were co­ administered with NGF at 5 min. Changes in fluorescence were calculated by subtracting the background from the fluorescence intensity at a specified time point ( F) and then normalized to the initial fluorescence intensity (F) resulting in ΔF/F.

### NGF-evoked nociception

Mouse NGF was administered by i.pl. injection into the right hindpaw of mice. EG00229, CendR or control peptides, compound 5 (36), a human monoclonal anti-NRP1 antibody (Vesencumab), or lgG control was administered (i.pl.) concomitantly with NGF.

### Inflammatory pain

CFA or vehicle (0.9% NaCl) was administered (i.pl.) into the right hindpaw of mice. NRP1 inhibitors were injected (i.pl.) 48 h after CFA.

### lntrathecal siRNA

Mouse GIPC1 or Ctrl siRNA was mixed with in vivo-jetPEI transfection reagent and administered to conscious mice (i.t., L4-L5), 48 h before NGF (i.pl.) or 48 h after CFA (i.pl.). *Gipc1* mRNA in DRG (L4-L5) was analyzed by RNAScope™ *in situ* hybridization, 48 h after siRNA injection.

### Nociception assays

Investigators were blinded to treatments. Mechanical allodynia and thermal hyperalgesia were measured (63).

### TrkA phosphorylation

Activated TrkA was detected in cultured DRG neurons by immunofluorescence using a rabbit mAb to phosphorylated TrkA Y785/TrkB Y816.

### ERK activity and ERK transcriptional activity

ERK activity was measured in cells expressing EKAR FRET biosensors. ERK transcriptional activity was measured in cells expressing SRE-Luc2P biosensors.

### ERK phosphorylation

Activated ERK and neurons were identified in cultured DRG neurons by immunofluorescence using a rabbit mAb to phosphorylated 44/42 (Thr202/Tyr204) and guinea pig polyclonal Anti-NeuN.

### Live cell imaging

Cells expressing SnapTag-TrkA and HaloTag-NRP1 were labeled with membrane­ impermeant SNAP-Surface® Alexa Fluor® 488 and membrane-impermeant HaloTag® Alexa Fluor® 660 and were imaged.

### Receptor-receptor BRET

For end point studies, HEK293T cells were transfected with a fixed concentration of Nanoluc-NRP1, Nanoluc-p75NTR or Nanoluc-TrkA and increasing concentrations of SnapTag-TrkA or SnapTag-CALCRL. For kinetic studies, cells were transfected with Nanoluc-TrkA, SnapTag-TrkA and either pcDNA3.1 or HaloTag-NRP1. Cells were incubated with SNAP-Surface® Alexa Fluor® 488. Luminescence and fluorescence were measured.

### Quantifying SNAP-Surface® Alexa Fluor® 488

HEK293T or CAD cells were transfected with SnapTag­ TrkA and pcDNA3.1 or HaloTag-NRP1. Cells were labeled using SNAP-Surface® Alexa Fluor® 488 and HaloTag® Alexa Fluor® 660, washed and fluorescence emissions were recorded.

### BRET trafficking

Cells were transfected with TrkA-HA-Rluc8 and fluorescent markers for the plasma membrane (RGFP-CAAX), early endosomes (tdRGFP-Rab5a), recycling endosomes (tdRGFP-Rab4a) or cis-Golgi apparatus (tdRGFP-Giantin). In some conditions, cells were co-transfected with pcDNA3.1 or HaloTag-NRP1, or siRNA. Cells were preincubated with hypertonic sucrose, clathrin inhibitor pitstop 2, GIPC1 inhibitor CR1023 or its negative control (CR2055), or myosin VI inhibitor TIP. Cells were incubated with luciferase substrate coelenterazine purple and stimulated with NGF. BRET ratios were recorded.

### Quantitative PCR (qPCR)

Primers to hTrkA (Hs01021011_m1), mTrkA (Mm01219406_m1), hNgfr (Hs00609976_m1), mNgfr (Mm00446296_m1), hGIPC1 (Hs00991802_m1), mGIPC1 (Mm00457561_m1), or GAPDH (Mn99999915_g1) were used. The relative abundance of mRNA was calculated (63). **Statistics.** Data are presented as mean ± SEM or ± SD. Differences were assessed using paired or unpaired t-test for two comparisons, and 1- or 2-way ANOVA and Tukey’s, Dunnett’s or Sidak’s post-hoc test for multiple comparisons. P<0.05 was considered significant at the 95% confidence level.

### Study approvals

Collection of DRG from deidentified organ donors was reviewed by the Institutional Review Board of the University of Cincinnati (#00003152, Study ID 2015-5302) and deemed to be human subjects exempt. New York University Institutional Animal Care and Use Committee approved experiments on animals (Protocol #PROTO202000006).

## Data availability

Primary data will be provided by the corresponding author upon request.

## Author Contributions

C.J.P., R.T., E.D., K.G., A.C-R, R.B., H.B., L.M. and A.M. conducted experiments and analyzed results; H.H., A.R.B.T. provided reagents; C.J.P., B.L.S., S.D., A.d.G., R.K. and N.W.B. designed experiments, wrote the manuscript and obtained funding.

## Acknowledgments

CJP was funded by the Leon Levy Fellowship in Neuroscience (CP, 2022-2023). Supported by grants from: National Institutes of Health (NS102722, DK118971, DE026806, DE029951, RM1DE033491 (NWB, BLS), GM147088 (ARBT), GM147088 (AdG), NS098772, NS120663, DA042852 **(RK),** NS134965 (KG) and Department of Defense (W81XWH1810431, W81XWH2210239, NWB, BLS). We thank Laura Kilpatrick and Stephen Hill from the University of Nottingham, and Promega Corporation for NRP1 cDNA plasmids. We thank Paz Duran for CAD cells.

