## Supplemental Material for "Neuropilin-1 is a co-receptor for Nerve Growth Factor-evoked pain"

### Materials and Methods

**Reagents.** Recombinant βNGF (R&D Systems or Alomone Labs) and recombinant NRP1 (R&D Systems) were reconstituted according to manufacturer's instructions. Unless stated otherwise, *in vitro* studies used human NGF, whereas ex vivo and in vivo studies used mouse NGF. EG00229 (HY-10799) was from MedChemExpress, compound 5 has been described (1), human monoclonal anti-NRP1 antibody (Vesencumab) was from Invitrogen (MA5-41940), and IgG control was from ThermoFisher (02-6502). Fragments of human NGF were from Genscript. The C-endR peptide includes the two conserved 'CendR' R/KxxR/K motifs within the C-terminus (underlined, QAAWRFIRIDTACVCVLSRKAVRRA) and corresponds to position 96-120 of mature NGF peptide. The control peptide (ARVAGQTRNITVDPRLFKKRRLRSP) corresponds to position 61-85 of immature NGF peptide. GIPC1 inhibitors (CR1023 or negative control) were from Genscript. Myosin VI inhibitor (2,4,6-triiodophenol) was from Thermo Scientific Chemicals.

**cDNAs.** SnapTag-TrkA was cloned with a GSSGAIA spacer and an N-terminal interleukin-6 signal sequence (IL-6-SnapTag-TrkA). IL-6-NanoLuc-TrkA, IL-6-NanoLuc-p75$^{NTR}$ and TrkA-HA-Rluc8 were cloned using Gibson assembly. HiBiT, a short fragment of NanoLuc (2), was cloned on the N-terminus of VEGF$_{165}$a or C-terminus of βNGF with a GSSG linker. HaloTag-NRP1, SnapTag-NRP1 and NanoLuc-NRP1 were from Dr. Laura Kilpatrick and Prof. Stephen Hill (University of Nottingham). SnapTag-CALCRL was from Alex Thomsen (New York University). RGFP-CAAX (prenylation CAAX box of KRas), tdRGFP-Rab5a, tdRGFP-Rab4a and tdRGFP-Giantin were from M. Bouvier (Université de Montréal). Nuclear and cytosolic EKAR (CFP/YFP) FRET biosensors were from Addgene.

**Cell Culture.** HEK293T cells were grown in complete Dulbecco's modified Eagle's medium GlutaMAX™ supplemented with 10% fetal bovine serum (FBS) and 100 U/mL penicillin-streptomycin. CAD cells were grown in RPMI 1640 containing 4% FBS, 8% horse serum and 100 U/mL penicillin-streptomycin.

**siRNAs.** ONTARGETplus siRNA targeting human GIPC1 (L-019997-00-0010), mouse GIPC1 (L-062534-00-0005) or nontargeting control siRNA (D-001810-10-05) were from Dharmacon (**Table S3**).

**Transfection of cell lines.** HEK293T cells were transfected using polyethylenimine (PEI, Polysciences; 1:6 DNA:PEI) diluted in a 150 mM NaCl solution. CAD cells were transfected with Lipofectamine 3000 (according to manufacturer's instructions) diluted in serum-free DMEM.

**Animals.** Experiments were in accordance with the guidelines recommended by the National Institute of Health, the International Association for the study of Pain, the National Centre for the Replacement, Refinement and Reduction of Animals in Research (ARRIVE) guidelines, and were approved by the New York University Institutional Animal Care and Use Committee (PROTO202000006). Male and female C57BL/6 mice (8-10 weeks, Jackson Laboratory) were housed four per cage at 22 ± 0.5°C under a controlled 14/10 h light/dark cycle with free access to food and water. Mice were randomly assigned to experimental groups; group size was based on previous similar studies. Investigators were blind to treatments.

**Microscale thermophoresis.** Human βNGF (100 µl, 10 µM; R&D Systems Inc.) was fluorescently labeled at lysine residues using the Monolith Protein Labeling Kit RED-NHS 2nd Generation (NanoTemper). Fluorescent NGF (15 nM final) was mixed with increasing concentrations of unlabeled NRP1-His (KactusBio) from 1.5 x $10^{-10}$ to 5 x $10^{-6}$ M in MST optimization buffer (50 mM Tris-Cl pH = 7.5, 150 mM NaCl, 10 mM $MgCl_2$, 0.05% Tween-20) and incubated at RT for 30 min. As a negative control, fluorescent NGF (15 nM final) was similarly incubated with staphylokinase from *Staphylococcus aureus* (Genscript). Samples were centrifuged (15,000 g, 10 min) and loaded on MST capillaries. Thermophoresis was measured using a Monolith NT.115 (NanoTemper Technology) at medium MST power. Data from four independent binding experiments was analyzed with GraphPad Prism using a non-linear regression fit (log[agonist] vs response).

**HiBiT-BRET binding assay**. HEK293T cells were transfected with SnapTag-NRP1 or SnapTag-NRP1 Y297A (3 µg/10 cm dish) using PEI. After 24 h, cells were transferred to 96-well plates. The next day, cells were incubated with SNAP-Surface® Alexa Fluor® 488 (0.25 µM, 1 h, 37ºC), then washed in HBSS/0.1% BSA. Separately, HEK293T cells were transfected with IL-6-HiBiT-VEGF$_{165}$a (3 µg/10 cm dish) or IL-6-NGF-HiBiT (5 µg/dish). After 48 h, medium was replaced with HBSS/0.1% BSA to collect secreted growth factor in cell supernatant (2 h, 37ºC). Supernatant was diluted (1:2) in HBSS/0.1% BSA and incubated with recombinant HaloTag-LgBiT protein and luciferase substrate furimazine (1:50 final dilution for both; Promega Corporation). NRP1-expressing cells were pre-incubated with vehicle or unlabeled VEGF$_{165}$a (10 nM, 30 min), followed by supernatant with furimazine (1:2, 15 min, 37ºC). Luminescence and fluorescence emissions were recorded (Synergy Neo2, Agilent BioTek; BRET1 donor 460 ± 40 nm, acceptor 540 ± 25 nm).

**Information-driven construction of a ternary NGF/TrkA/NRP1 computational model.** The TrkA extracellular domain (ECD) structure from the crystal structure of the NGF/TrkA complex (PDB 2IFG) and the AlphaFold (3) -derived model of hNRP1 with only a1a2b1b2 domains were used in docking calculations. hNGF has two prospective 'CendR' R/KxxR/K motifs within the C-terminus that are important for mediating interactions of other growth factors with NRP1 b1 domains (4, 5). The AlphaFold-derived model of soluble hNGF sterically clashed with TrkA when superimposed with the NGF/TrkA crystal structure. Further, the NGF C-terminus CendR$_{118}$ motifs were missing density in NGF/TrkA crystal structure. Hence, Modeller (6) was used to add missing structure to NGF in the NGF/TrkA crystal structure and to sample conformations of NGF CendR$_{118}$ motifs for docking calculations. When modeled on the cell membrane, analysis of the NGF/TrkA crystal structure (PDB 2IFG) revealed that hNGF CendR motif R$_{100}$FIR$_{103}$ is membrane proximal whereas motif K$_{115}$AVR$_{118}$ is membrane distal. hNGF polarity with respect to the membrane was determined by the relative membrane orientation of bound hTrkA ECDs (PDB 2IFG). This analysis identified two probable binding modes of NRP1 to the membrane-tethered NGF/TrkA complex. In mode 1, NRP1 b1 domain is poised to interact with CendR motif R$_{100}$FIR$_{103}$. In this mode, a1a2b2 domains are membrane proximal and NRP1 can possibly interact with the membrane. In mode 2, NRP1 b1 domain is poised to interact with CendR motif R$_{115}$FIR$_{118}$. This time, the a1a2b2 domains are membrane distal and NRP1 can also possibly interact with the TrkA ECD domains. However, in mode 1, the NGF CendR motif R$_{100}$FIR$_{103}$ is at the interface with TrkA Ig-C2 domain and is not accessible for NRP1 binding. Hence, we computationally explored binding mode 2 and analyzed its relevance in a physiological context as outlined below.

The NGF/TrkA/NRP1 model was generated in a data-driven approach using HADDOCK 2.4 webserver (7). Since TrkA binding to NGF does not utilize CendR motif R$_{115}$FIR$_{118}$ of NGF, NRP1 was directly docked to the NGF/TrkA crystal structure. NGF models with solvent-accessible conformations of CendR motif R$_{115}$FIR$_{118}$ in the NGF/TrkA complex were selected as first docking partners and the AlphaFold-derived NRP1 a1a2b1b2 domain structure as the second docking partner. Knowledge-based restraints in the form of active and passive residues were used as restraints in the docking protocol. In HADDOCK, active residues are solvent-exposed residues directly involved in the interaction between the two proteins followed by solvent-exposed passive residues close to the active residues that may be at the interface (8). From NGF/TrkA, residues of CendR motif R$_{115}$FIR$_{118}$ and from NRP1, b1 domain residues Y297, D320, S346, T349 and Y353 were specified as active residues. These NRP1 b1 domain residues constitute a C-terminal arginine binding pocket that is known to bind C-terminal arginine in NRP-binding proteins and peptides (4, 5, 9, 10). All residues within 6.5 Å of active residues were specified as passive residues in both docking partners. Standard docking parameters were used during the rigid-body energy minimization followed by semi-flexible refinement where residues at the interface are allowed to move. Besides docking score, several factors were considered while selecting the docked models. This was to ensure the selection of sterically feasible binding modes of NRP1 to NGF/TrkA complex modeled on the membrane without clashing with either the membrane or TrkA. Although non-standard residues were excluded from docking calculations, compatibility of glycosylated TrkA with NRP1 binding in cellular context was considered while selecting docked models (**Fig. S1A**). Further, suitable orientations of the NRP1 b2 domains in connection with NRP1 MAM domain were also considered during selection to ensure membrane-tethered NGF/TrkA/NRP1 complex formation were also possible (**Fig. 1F**). In the different docked poses derived from docking calculations, the NRP1 b1 domain still binds to the NGF C-terminus in a similar manner while

NRP1 a1, a2 and b2 domains are oriented differently resulting in poses where one NRP1 molecule can bind to one TrkA monomer (**Fig. 1E**) or two TrkA monomers at once thereby bridging TrkA dimers (**Fig. S1B**). Explicit water refinement was then performed on the selected docked models using the refinement interface of HADDOCK 2.4 webserver (7). The refined ternary complexes were analyzed for molecular interactions at the interfaces using PISA (11). Structural superposition of structural complexes for analysis and preparation of figures was made using PyMol (12) and UCSF ChimeraX (13).

**Collection of mouse tissue.** Mice were anesthetized (5% isoflurane) and perfused through the ascending aorta with PBS and then 4% paraformaldehyde in PBS. DRG (L4-L5) were removed, fixed in 4% paraformaldehyde in PBS (1 h, 4°C), cryoprotected in 30% sucrose (24 h, 4°C), and embedded in Optimal Cutting Temperature compound (Tissue Tek). Frozen sections (10-12 µm) were mounted, dried (15 min) and stored (-20°C).

**Collection of human tissue.** The collection of DRG from deidentified organ donors was reviewed by the Institutional Review Board of the University of Cincinnati (#00003152, Study ID 2015-5302) and deemed to be human subjects exempt. Donor information has been provided (14). DRG (L4-L5) were collected in the operating room within 90 min of aortic cross clamp and after removal of vital organs. DRG were placed in N-methyl-D-glucamine-artificial cerebrospinal fluid (4°C), dissected and immersion fixed in 4% paraformaldehyde in PBS (overnight, 4°C), cryoprotected in 30% sucrose (24 h, 4°C), and embedded in Optimal Cutting Temperature compound. Frozen sections (14-18 µm) were mounted, dried (15 min) and stored (-80°C).

**Immunofluorescence.** Mouse tissue was blocked (2% bovine serum albumin, BSA, 0.2% Triton X-100 in PBS; 1 h, RT). Sections were incubated (4ºC, overnight) with monoclonal rabbit anti-TrkA (1:50; Millipore-Sigma, SU0354) or polyclonal rabbit anti-glutamine synthetase antibody (GS, 1:15000, Abcam, ab49873), and monoclonal mouse anti-NRP1 (1:200; Santa Cruz, A-12) or monoclonal human anti-NRP1 Vesencumab (1:500; Invitrogen, MA5-41940). Slides were washed and incubated with donkey anti-rabbit Alexa Fluor® 488 (1:1000) and either donkey anti-mouse or anti-human Alexa Fluor® 647 (1:1000, 1 h, RT). Slides were incubated with DAPI (1 µg/ml, 5 min) and mounted (ProLong® Gold Antifade). Sections were imaged using a Leica SP8 confocal microscope with HCX PL APO 40x oil objective (Leica Microsystems).

**RNAScope™ *in situ* hybridization in mouse and human tissue.** mRNA transcripts were localized in mouse and human tissues using a RNAScope™ Multiplex Fluorescent Reagent v2 Assay Kit (Advanced Cell Diagnostics Inc.) using the protocol for fresh-frozen tissue as recommended by the manufacturer, except for omission of the initial on-slide fixation step with mouse tissue. Probes to mouse (Mn) Mm-Nrp1(#471621-C3), Mm-Ntrk1 (#435791-C2), Mm-Gipc1 (#1232971-C1) and human (Hs) Hs-Nrp1 (#424361-C1), Hs-Ntrk1 (#402631-C2) and Hs-Gipc1 (#1232961-C3) were used. Mouse tissue was incubated with Opal 620 (1:1000, FP1495001KT, Akoya Biosciences) and Opal 520 (FP1487001KT) reagents for fluorescence detection. For human tissue, Tyramide Signal Amplification Plus Cyanine 3 (1:1500) and Cyanine 5 (1:1000) were used to detect the hybridized signals. To detect all neurons or peptidergic neurons in mouse tissues, hybridized slides were blocked and incubated with guinea pig anti-NeuN antibody (1:500, EMD Millipore, N90) or rabbit anti-CGRP (1:1000, cat#C8198, Sigma), respectively (overnight, 4°C). Slides were washed and incubated with goat anti-guinea pig Alexa Fluor® 647 (1:1000) or goat anti-rabbit Alexa Fluor® 488 (1:1000, ThermoFisher) (1 hour, RT). Alternatively, neurons were detected using Nissl staining. Slides were washed, incubated with DAPI and mounted. For mouse tissue, sections were imaged using a Leica SP8 confocal microscope with HCX PL APO 40x (NA 1.30) oil objective. The percentage of hybridized positive neurons were quantified and normalized by the total number of neurons. The percentage of hybridized positive peptidergic neurons were quantified and normalized by the total number of peptidergic neurons. To detect peptidergic and non-peptidergic neurons in human tissues, hybridized slides were blocked and incubated with mouse anti-CGRP (1:500, cat# C7113, Sigma) or rabbit anti-P2X3 (1:500, cat# NBP2-33848, Novus Biologicals) respectively (overnight, 4°C). Slides were washed and incubated with donkey anti-mouse Alexa Fluor® 488 or donkey anti-rabbit Alexa Fluor® 555 (1:1000, ThermoFisher) (1 hour, RT). For human tissue, images were captured as stitched Z stacks using Keyence BZ-X800E fluorescence microscope. Cells positive for *Ntrk1*, *Nrp1* and *Gipc1* were counted to mark positive cells, where 10+ fluorescent puncta were considered to be positive.

**Mouse DRG cultures for calcium imaging.** C57BL/6 mice (~4-6 weeks) were anaesthetized (5% isoflurane). All DRGs were excised and dissociated with collagenase type I (1 mg/ml; 30 min, 37°C), followed by papain (0.125 mg/ml; 30 min, 37°C). Dissociated cells were pelleted and resuspended in DMEM containing 10% FBS and 1% penicillin-streptomycin. Neurons were plated on MatTEK 35 mm dishes (P35G-1.5-14-C) coated with poly-D-lysine (0.1 mg/ml; 20 min; Sigma-Aldrich; P7280) and imaged within 24 h.

**Calcium imaging on mouse DRG neurons**. DRG neurons were incubated with Fluo-4AM (1 µM; Tocris; 6255) in a calcium buffer (in mM: 150 NaCl, 2.5 KCl, 2.2 $CaCl_2.2H_20$, 1.18 $MgCl_2.6H_2O$, 10 D-Glucose, 10 HEPES, 0.5% BSA, 1X Probenecid, pH 7.4; 1 h at 37°C). Neurons were continuously perfused (10 ml/min) with calcium buffer and maintained at 37°C using the Tokai Hit incubation system (ZILCS S/N 172288). Fluorescence was recorded in individual neurons using a Leica DMi8 microscope equipped with a HC PL FLUOTAE 10x (NA 0.30) air objective (Leica Microsystem) and a Leica DFC9000GTC camera. Images were collected every 5 s to minimize photobleaching and phototoxicity. After basal recording for 1 min, neurons were challenged with capsaicin (100 nM) at 1 min and at 7 min. Depolarization was evoked with 50 mM KCl at 13 min. Neurons were exposed to mouse NGF (100 nM) or vehicle (control) at 5 min. In some experiments, neurons were preincubated with EG00229 (3, 10 or 30 µM) or vehicle (0.1% DMSO, control) for 1 min before capsaicin. CendR or control peptide (0.1, 0.3 or 1 µM) and Vesencumab mAb or control IgG (0.7 µg/ml) were co-administered with NGF at 5 min. Changes in fluorescence were calculated by subtracting the background from the fluorescence intensity at a specified time point (ΔF) and then normalized to the initial fluorescence intensity (F) resulting in ΔF/F.

**Human DRG cultures.** Human DRG suspension cells were from AnaBios Corporation. Cells were recovered with a gentle centrifugation (RT, 3 min) and resuspension with DMEM/F12 medium containing 10% horse serum, 1% penicillin-streptomycin, 25 ng/ml hNGF and 25 ng/ml GDNF. Cells were seeded on poly-D-lysine (0.1 mg/ml) and laminin (1 mg/ml)-coated 12-mm glass coverslips. Half of the culture medium was replaced with fresh medium every 3 days. Cultures used within 96 h.

**Mouse DRG cultures for electrophysiology.** C57BL/6 mice (~3-4 weeks) were anaesthetized (5% isoflurane). Thoracic and lumbar DRG were excised and dissociated with collagenase type I (1 mg/ml) and neutral protease (0.62 mg/ml; 45 min, 37°C) under gentle agitation. Dissociated cells were pelleted and resuspended in DMEM containing 10% FBS and 1% penicillin-streptomycin. For siRNA transfection, cells were resuspended in Nucleofector® transfection reagent containing GIPC1 or control siRNA (500 nM) and a plasmid encoding the enhanced green fluorescent protein (eGFP, 1 µg). Cells were electroporated using Amaxa Biosystem (Lonza) with protocol O-003. Neurons were plated on glass coverslips pretreated with poly-D-lysine (0.1 mg/ml) and laminin (0.01 mg/ml; 30 min). Neurons were used within 48 h and siRNA transfection verified by eGFP fluorescence.

**Whole-cell patch-clamp recordings of action potentials and rheobase.** DRG neurons were preincubated with EG00229 (10 or 30 µM, 30 min) or vehicle (0.1% DMSO) and then challenged with mouse or human NGF (50 nM, 30 min). For current-clamp recordings, the external solution contained (mM): 154 NaCl, 5.6 KCl, 2 $CaCl_2$, 1 $MgCl_2$, 10 D-Glucose, and 8 HEPES (pH 7.4 adjusted with KOH, and mOsm/L= 300). The internal solution comprised (mM): 137 KCl, 10 NaCl, 1 $MgCl_2$, 1 EGTA, and 10 HEPES (pH 7.3 adjusted with KOH, and mOsm/L= 277). Recordings of action potentials were made at RT in whole-cell patch clamp configuration and current-clamp mode. DRG neurons with a resting membrane potential (RMP) more hyperpolarized than −40 mV, stable baseline recordings, and evoked spikes that overshot 0 mV were used for experiments and analysis. Action potentials were evoked by a ramp pulse from 0-1000 pA in 1 sec for human DRG, or 0-250 pA in 1 sec for mouse DRG. Rheobase was determined from the current ramp pulses in both human and mouse conditions. Results were analyzed using Fitmaster software (HEKA), Easy Electrophysiology 2.7.3 and Origin 9.0 software (OriginLab).

**Whole-cell patch-clamp recordings of $Ca^{2+}$ and $Na^+$ currents.** DRG neurons were preincubated with EG00229 (30 µM, 30 min) or vehicle (0.1% DMSO) and then challenged with mouse or human NGF (50 nM, 30 min). $Ca^{2+}$ currents ($ICa^{2+}$) and $Na^+$ ($INa^+$) currents were recorded as described (15, 16). In brief, peak $Ca^{2+}$ current was acquired by applying 200 msec voltage steps from −70 to +70 mV in 10 mV increments from a holding potential of −90 mV to obtain the current-voltage (I-V) relation. Peak $Na^+$ current was acquired by applying 150 msec voltage steps from −70 to +60 mV in 5 mV increments from a holding potential of −60 mV to obtain the current-voltage (I-V) relation. Capacitive artifacts were fully compensated, and series resistance was compensated by ~70 %. Recordings made from cells with greater than a 20% shift in series resistance compensation error were excluded from the analysis. All recordings were made at RT (~23 °C). Results were analyzed with Patchmaster software.

**NGF-evoked nociception.** Mouse NGF (50 ng/10 µl) or vehicle was administered by intraplantar (i.pl.) injection into the right hindpaw of mice. EG00229 (1, 10 or 30 µM/10 µl), compound 5 (30 µM/10 µl) (1), CendR or control peptide (0.2, 2 or 10 µM/10 µl), human monoclonal anti-NRP1 antibody (Vesencumab) or IgG control (7 µg/10 µl) was administered (i.pl.) concomitantly with NGF. GIPC1 or control siRNA was administrated by intrathecal (i.t) injection 48 h before NGF (i.pl.).

**Inflammatory pain.** CFA (1 mg/ml) or vehicle (0.9% NaCl) was administered (10 µl, i.pl.) into the right hindpaw of sedated mice (2% isoflurane). NRP1 inhibitors were injected (i.pl.) 48 h after CFA. GIPC1 or control siRNA was administrated (i.t) 24 h after CFA.

**Intrathecal administration of siRNA.** Mouse GIPC1 or control siRNA (1.25 µg) was mixed with in vivo-jetPEI transfection reagent (8:1 PEI:DNA; Polyplus, 201-50G) and administered to conscious mice (5 µl, i.t., L4-L5), 48 h before NGF (i.pl.) or 24 h after CFA (i.pl.). *Gipc1* mRNA in DRG (L4-L5) was analyzed by RNAScope$^{TM}$ *in situ* hybridization, 48 h after siRNA injection.

**Nociception assays.** Investigators were blinded to treatments. Mechanical allodynia was assessed by measuring hindpaw withdrawal response to VFF stimulation using the up-and-down method (14). The Hargreaves apparatus was used to evaluate hypersensitivity to heat (Ugo Basile) (14).

**Phosphorylated TrkA assays.** Acutely dispersed mouse DRG neurons were plated onto coverslips pre-coated with Cell-Tak (30 min, 37°C; CB-40240) in DMEM containing 10% FBS. After 18 h, neurons were incubated with a monoclonal NGF antibody (1:500, 4 h, 37°C; MA5-41968) in serum-free DMEM to reduce interference from endogenous NGF. Neurons were washed in HBSS/0.1% BSA, pre-incubated with EG00229 (30 µM, 30 min) or control, then stimulated with vehicle or mouse NGF (100 nM, 15 min, 37°C). Neurons were fixed with 4% paraformaldehyde in PBS (20 min, 4°C), blocked (2% BSA, 0.2% Triton-X-100 in PBS), and incubated with a rabbit mAb to phosphorylated TrkA Y785/TrkB Y816 (1:500, overnight, 4°C; Cell Signaling, 4168). Neurons were washed in PBS and incubated with donkey anti-rabbit Alexa Fluor® 488 (1:1,000, 1 h, RT). Slides were incubated with DAPI, mounted then imaged using a Leica SP8 confocal microscope with HCX PL APO 40x (NA 1.30) oil objective. Mean fluorescence intensity was quantified from regions of interest drawn using a phase contrast image with the maximum projection from four Z planes (0.5 µm thickness). Images were quantified from 34-44 individual cells across three independent experiments using separate mice.

**ERK activity FRET biosensor assays.** CAD cells were serum-starved and transfected with EKAR FRET biosensors (300 ng/well) and human SnapTag-TrkA (150 ng/well). HEK293T-FLPN cells were transfected with FRET biosensors (1 µg/10 cm dish), SnapTag-TrkA (1 µg), and pcDNA3.1 or HaloTag-NRP1 (2 µg). After 24 h, cells were transferred to 96-well plates and serum-starved overnight. On the day of the assay, cells were washed and incubated in Hanks' buffered saline solution (HBSS, pH 7.4) containing 0.1% BSA. After 30 min (37°C), FRET was measured at 60 s intervals (CLARIOstar, BMG Labtech). After 5 baseline reads, cells were stimulated with NGF (0.1 pM-10 nM) or phorbol 12,13-dibutyrate (PDBu, 10 µM). ΔFRET represents the ratio (YFP/CFP), minus the mean from 5 initial reads and baseline-corrected to vehicle. Area under the curve (AUC) was determined for each replicate.

**ERK transcriptional assays.** HEK293T-FLPN cells were transfected with SRE-Luc2P (3 µg/10 cm dish), SnapTag-TrkA (1 µg) and either pcDNA3.1 or HaloTag-NRP1 (2 µg). After 24 h, cells were transferred to 96-well plates. CAD cells were transfected with SRE-Luc2P (250 ng/well, 96-well plate), SnapTag-TrkA (150 ng/well) and 100 nM siRNA. After 24 h, cells were serum-starved overnight. Cells were then incubated in HBSS/0.1% BSA and stimulated with NGF (1 pM-10 nM) or PDBu (10 µM). After 5 h stimulation (37ºC), cells were incubated with luciferin (1:2) and lysis buffer (1:5). After 5 min to enable the substrate reaction, luminescence emissions were recorded (CLARIOstar).

**Phosphorylated ERK assays.** Mouse DRG neurons were plated onto coverslips pre-coated with poly-D-lysine (0.1 mg/ml; 20 min) in DMEM containing 10% FBS. After 18 h, neurons were incubated with a monoclonal NGF antibody (1:500, 4 h, 37°C; MA5-41968) in serum-free DMEM. Neurons were incubated with EG00229 (30 µM), CendR (1 µM), or controls, and vehicle or mouse NGF (100 nM) for 30 min at 37°C. Neurons were washed with PBS and fixed with 4% paraformaldehyde in PBS (15 min, 4°C), blocked (10%

normal donkey serum in PBS) for 1 h, and incubated with a rabbit mAb to phosphorylated 44/42 Thr202/Tyr204 ERK1/2 (1:400; Cell Signaling; 4168) and a guinea pig polyclonal anti-NeuN (1:800; Sigma-Aldrich; ABN90) in blocking solution for 2 h at RT. Neurons were washed in PBS and incubated with donkey anti-guinea pig Alexa Fluor® 488 (1:1,000) and donkey anti-rabbit Alexa Fluor® 568 (1:1,000, 45 min, RT). Slides were incubated with DAPI, mounted, and stored at 4°C. Neurons were imaged using a Leica SP8 confocal microscope with HC PL APO 20x (NA 0.75) air objective (Leica Microsystem), employing identical illumination exposure parameters for all groups. The FIJI-Image J cell counter plug-in was utilized to count the number of NeuN-positive cells and p-ERK/NeuN double-positive cells. Images were quantified from 15-432 individual cells across 4 independent experiments using separate mice.

**Live imaging.** HEK293T cells were transfected with SnapTag-TrkA (1 µg/10cm dish), HaloTag-NRP1 (2 µg) or both TrkA and NRP1. After 24 h, cells were transferred to 35 mm dishes. CAD cells were seeded in 35 mm dishes, serum-starved for 24 h, and then transfected with SnapTag-TrkA (250 µg/35 mm), HaloTag-NRP1 (500 µg) or both. The next day, proteins were labeled with SNAP-Surface® Alexa Fluor® 488 and membrane-impermeant HaloTag® Alexa Fluor® 660 (0.5 µM, 30 min, 37°C). Cells were washed in HBSS and imaged on the Leica SP8 confocal.

**Receptor-receptor BRET**. For endpoint studies, HEK293T cells were transfected with a fixed concentration of NanoLuc-NRP1, NanoLuc-p75$^{NTR}$ or NanoLuc-TrkA (10 ng/well, 96-well plate) and increasing concentrations of SnapTag-TrkA or SnapTag-CALCRL (0-50 ng/well). For kinetic studies, cells were transfected with NanoLuc-TrkA (10 ng/well), SnapTag-TrkA (25 ng/well) and either pcDNA3.1 or HaloTag-NRP1 (50 ng/well). After 48 h, cells were incubated with SNAP-Surface® Alexa Fluor® 488 (0.5 µM, 30 min, 37°C), washed in HBSS before fluorescence emissions were measured (Synergy Neo2). In some conditions, cells were incubated with NGF (30 nM, 30 min, 37°C) or vehicle. Cells were then incubated with furimazine (10 µM, 5 min, Promega). For kinetic studies, cells were stimulated with increasing NGF concentrations (1 pM-100 nM) following 5 baseline reads. Luminescence and fluorescence emissions were quantified (Synergy Neo2; BRET1 filter).

**Quantifying SNAP-Surface® Alexa Fluor® 488**. HEK293T cells were transfected with SnapTag-TrkA (25 ng/well, 96-well plate) and pcDNA3.1 or HaloTag-NRP1 (50 ng/well). CAD cells were transfected with SnapTag-TrkA (150 ng/well) and either pcDNA3.1 or HaloTag-NRP1 (300 ng/well). Cells were labeled using SNAP-Surface® Alexa Fluor® 488 and HaloTag® Alexa Fluor® 660 (0.5 µM, 30 min, 37°C), washed in HBSS/0.1% BSA and fluorescence emissions were recorded (CLARIOstar).

**BRET trafficking.** HEK293T cells were transfected with TrkA-HA-Rluc8 (15 ng/well, 96-well plate) and a fluorescent marker (15 ng/well) for the plasma membrane (RGFP-CAAX), early endosome (tdRGFP-Rab5a), recycling endosome (tdRGFP-Rab4a) or cis-Golgi apparatus (tdRGFP-Giantin). In some conditions, cells were co-transfected with pcDNA3.1 or HaloTag-NRP1 (30 ng/well), or 100 nM siRNA. Alternatively, CAD cells were transfected with TrkA-HA-Rluc8 (125 ng/well) in the absence or presence of HaloTag-NRP1 (250 ng/well), as well as RGFP-CAAX, tdRGFP-Rab5a, tdRGFP-Rab4a or tdRGFP-Giantin (150 ng/well). After 48 h, cells washed in HBSS/0.1% BSA were pre-incubated (30 min, 37°C) with hypertonic sucrose (0.45 M), Pitstop 2 (30 µM; Abcam), an inhibitor of GIPC1 (CR1023, 300 µM) or its negative control (CR2055), or an inhibitor of myosin VI (TIP, 50 µM). Cells were incubated with coelenterazine purple (25 µg/ml, 10 min, Nanolight). For kinetic studies, 5 baseline reads were measured, then cells were stimulated with NGF (1 pM-100 nM), using 30 nM NGF for endocytic inhibitors. BRET ratios were recorded every 45 s (Synergy Neo2; BRET2 filter: donor 410 ± 80 nm, acceptor 515 ± 30 nm). ΔBRET represents BRET in the presence of agonist, minus the vehicle BRET over time. For endpoint studies lacking agonist stimulation, BRET ratios were baseline-corrected to TrkA-HA-Rluc8 expression alone (100%).

**Quantitative PCR (qPCR).** RNA was isolated from cells or snap-frozen tissues using Direct-zol RNA MiniPrep Kit (Zymo Research, R2050). cDNA was prepared using MultiScribe Reverse Transcriptase (Thermo Fisher, 4311235). cDNA was amplified for 40 cycles by qRT-PCR using the QuantStudio 3 Real-Time PCR System and TaqMan Fast Advanced PCR Mastermix (4444556). Primers to hTrkA (Hs01021011_m1), mTrkA (Mm01219406_m1), hNgfr (Hs00609976_m1), mNgfr (Mm00446296_m1), hGIPC1 (Hs00991802_m1), mGIPC1 (Mm00457561_m1), or GAPDH (Mn99999915_g1) were used. The relative abundance of mRNA was calculated as described (14).

**Data analysis and statistics**. Images processed using ImageJ. Illustrations were produced using Adobe Illustrator, Mol*Viewer or BioRender. Data are presented as mean ± SEM, determined using GraphPad Prism (8.0). Concentration-response data were fitted using non-linear regression analysis as log *vs.* response (three parameters) to determine $EC_{50}$ values. Differences were assessed using paired or unpaired t-test for two comparisons, and 1- or 2-way ANOVA and Tukey's, Dunnett's or Šidák's post-hoc test for multiple comparisons. $P<0.05$ was considered significant at the 95% confidence level.

**SI Figures**

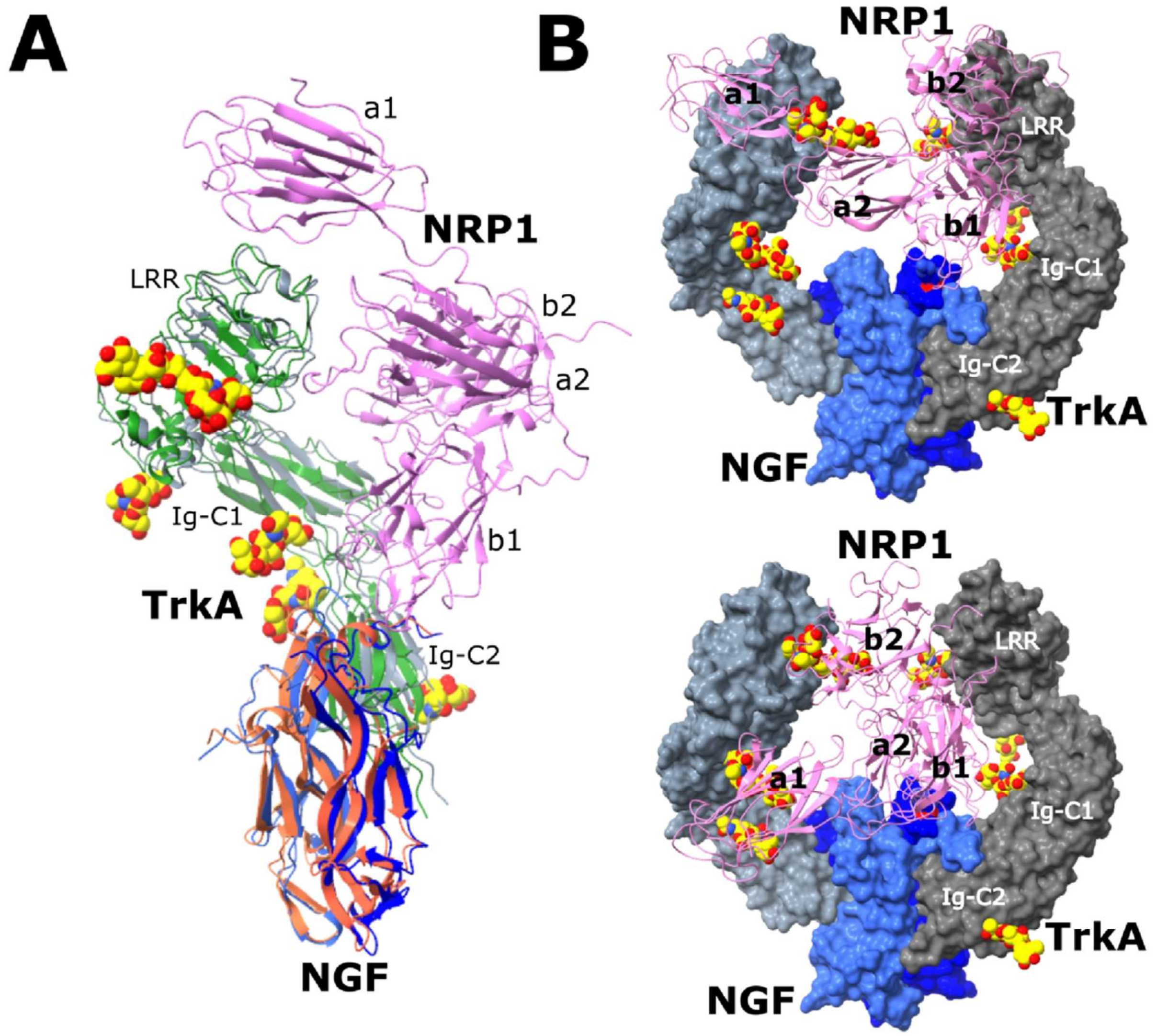

**Figure S1. NGF/TrkA/NRP1 docked models in the context of glycosylated TrkA extracellular domains. A.** Side view of structural superposition of NGF/TrkA/NRP1 (cartoon, blue/grey/pink) docked model and NGF/TrkA (cartoon, orange/green) crystal structure (PDB 2IFG) through common NGF/TrkA component is shown (only monomers are shown for NRP1 and TrkA for clarity). Asn-linked glycosylations on TrkA in NGF/TrkA crystal structure are shown as space-filling representation and colored according to atom type (C/O/N, yellow/red/blue). Individual domains are labeled and described in text. In cellular context, this comparison shows a glycosylated TrkA compatible binding mode of NRP1 in the docked NGF/TrkA/NRP1 model. **B.** Other docked poses of NRP1 observed in NGF/TrkA/NRP1 docking calculations are shown. Only one monomer of NRP1 (cartoon, pink) bound to NGF/TrkA (surface, blue/grey shades) is shown. For context, TrkA Asn-linked glycosylations are shown in space-filling representations. Individual domains are labeled and described in the text. In contrast to NGF/TrkA/NRP1 docked model described in text, one NRP1 molecule binds to two TrkA monomers at once thereby bridging TrkA dimers in these docked poses. Although NRP1 domains appear to interfere with TrkA glycosylations, domain flexibility in the cellular context (not accounted for in docking calculations) may allow these scenarios to exist. It is important to note that in different docked poses from docking calculations, NRP1 b1 domain still binds to NGF C-terminus in a similar manner while NRP1 a1, a2 and b2 domains are oriented differently.

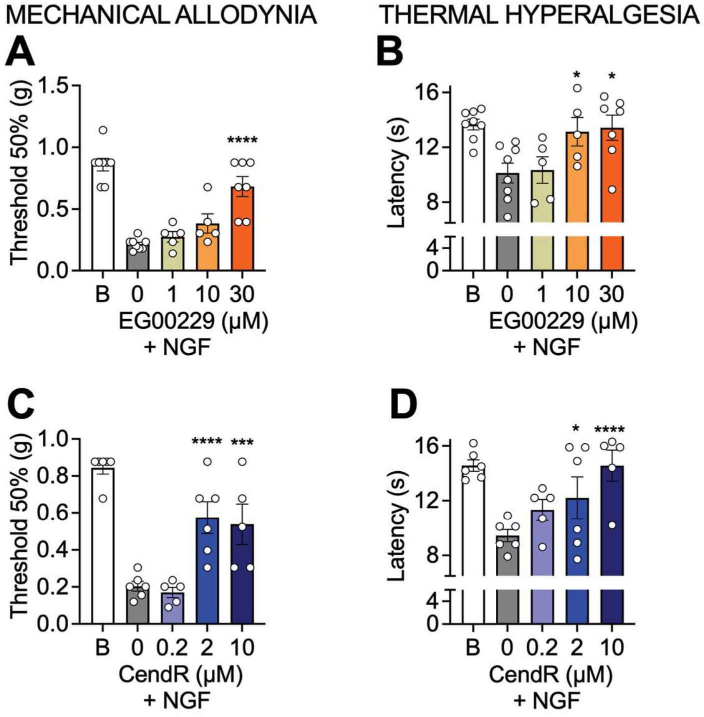

**Figure S2. Effect of Nrp1 inhibitors on NGF-induced nociception in mice.** Dose-response curve of NRP1 inhibitors EG00229 (**B, E**; 1, 10 and 30 µM/10 µl i.pl.) and CendR (**C, F**; 0.2, 2 and 10 µM/10 µl i.pl.). After baseline (B) measurements, inhibitors were co-injected with mouse NGF (50 ng/10 µl, i.pl.). Mechanical allodynia (A-C) and thermal hyperalgesia (B-D) were measured 1 hour after injection. N=5-8 male mice per group. Mean±SEM. **P*<0.05, ****P*<0.001, *****P*<0.0001 vs. 0 µM. 1-way ANOVA, Dunnett multiple comparisons.

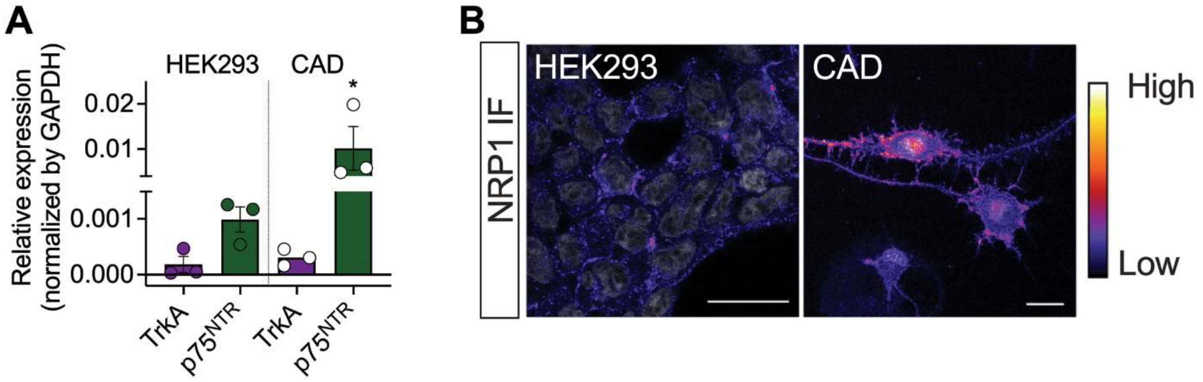

**Figure S3. NGF receptor expression in HEK293 and neuron-like CAD cells**. **A.** Quantification of expression of *Ntrk1* mRNA (TrkA) and *Ngfr* ($p75^{NTR}$) in HEK293T or CAD cells determined by quantitative RT-PCR. Data from 3 independent experiments with triplicate wells. **B.** Immunofluorescence (IF) staining of NRP1 in HEK293T and CAD cells using vesencumab. Scale bar, 20 µm. Representative image from 4 independent experiments. Mean±SEM. *$P$<0.05. 1-way ANOVA, Dunnett's multiple comparisons.

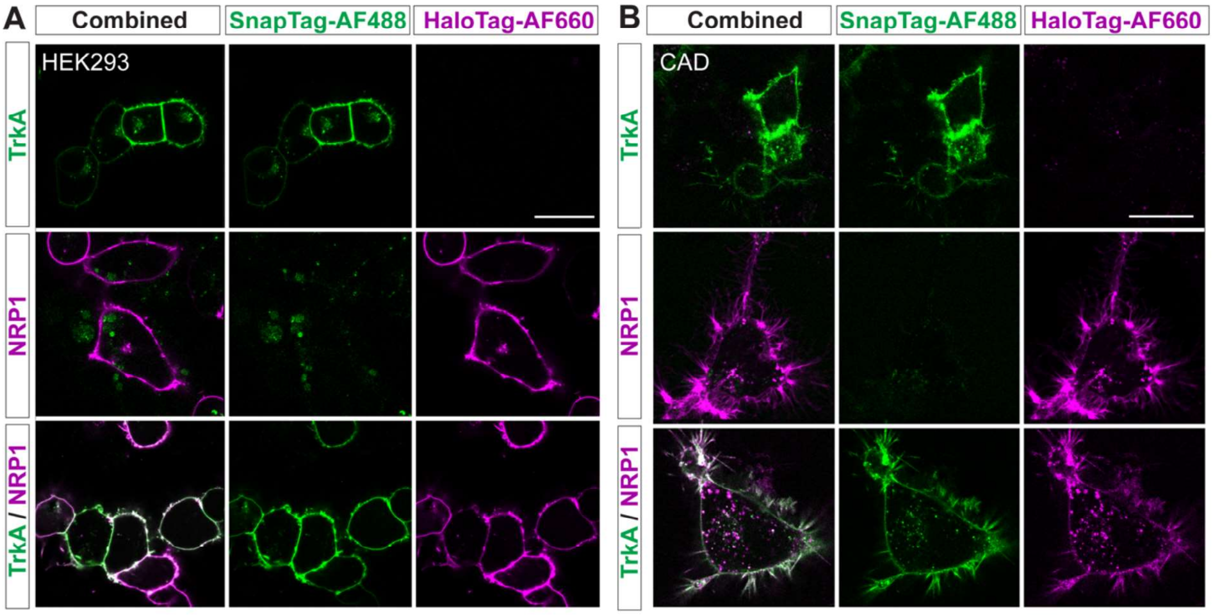

**Figure S4. Specificity of SnapTag- and HaloTag- labeling in live cells.** HEK293T cells (**A**) or CAD cells (**B**) were transfected with either SnapTag-TrkA or HaloTag-NRP1 or both (1:2 TrkA to NRP1 ratio). Tags were simultaneously labeled with membrane-impermeant substrates (SNAPTag-Alexa Fluor® 488, HaloTag-Alexa Fluor® 660). Representative from N=5 independent experiments. Scale bar, 20 μm.

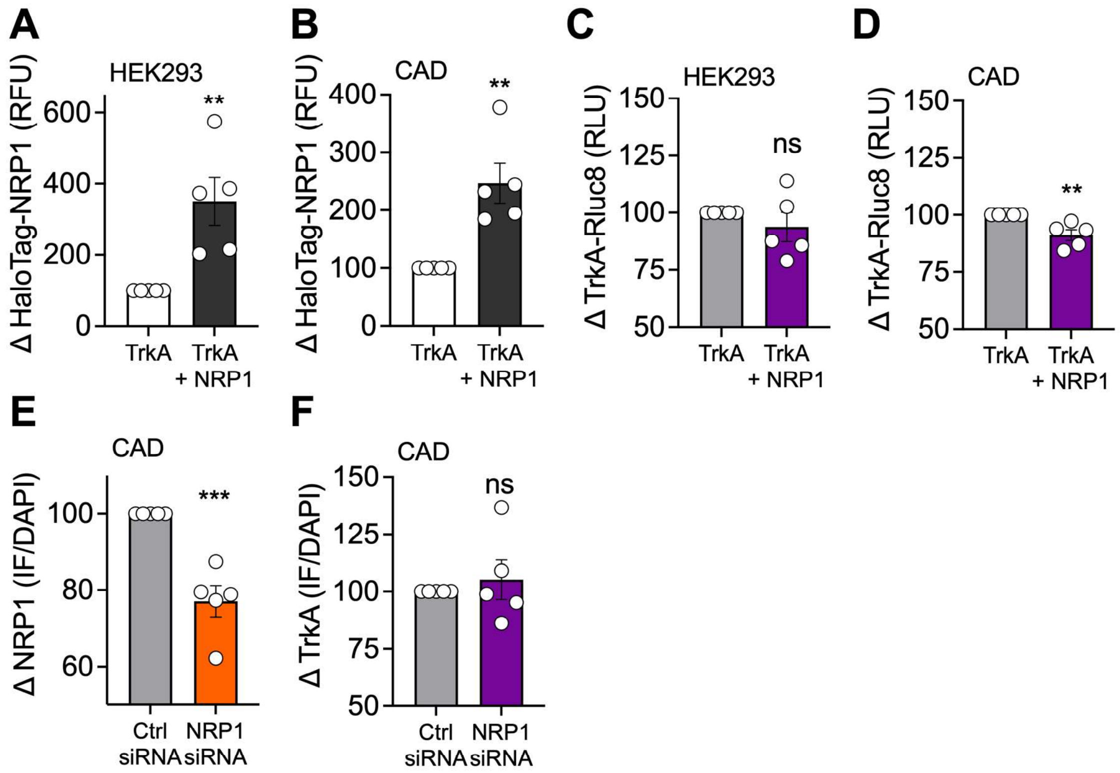

**Figure S5. Effect of co-expression or siRNA on TrkA and NRP1 expression in HEK293 and CAD. A cells. A-D.** NRP1 and TrkA expression in HEK293T (**A, C**) or CAD (**B, D**) cells expressing TrkA-Rluc8 in the absence or presence of HaloTag-NRP1. **A, B.** Cells were labeled with membrane-impermeant HaloTag-Alexa Fluor® 660 and emissions were measured as relative fluorescence units (RFU). **C, D.** TrkA-Rluc8 emissions were measured after incubation with coelenterazine purple to infer TrkA expression as relative luminescence units (RLU). **E, F.** Effect of NRP1 siRNA on expression of NRP1 (**E**) or TrkA (**F**) in CAD cells, quantified by immunofluorescence (IF) staining normalized to nuclei count (DAPI). Data expressed as percentage change over control (100%). Data from 5 independent experiments. Mean±SEM. ***P*<0.01, *** *P*<0.001, ns, not significant. Unpaired t-test.

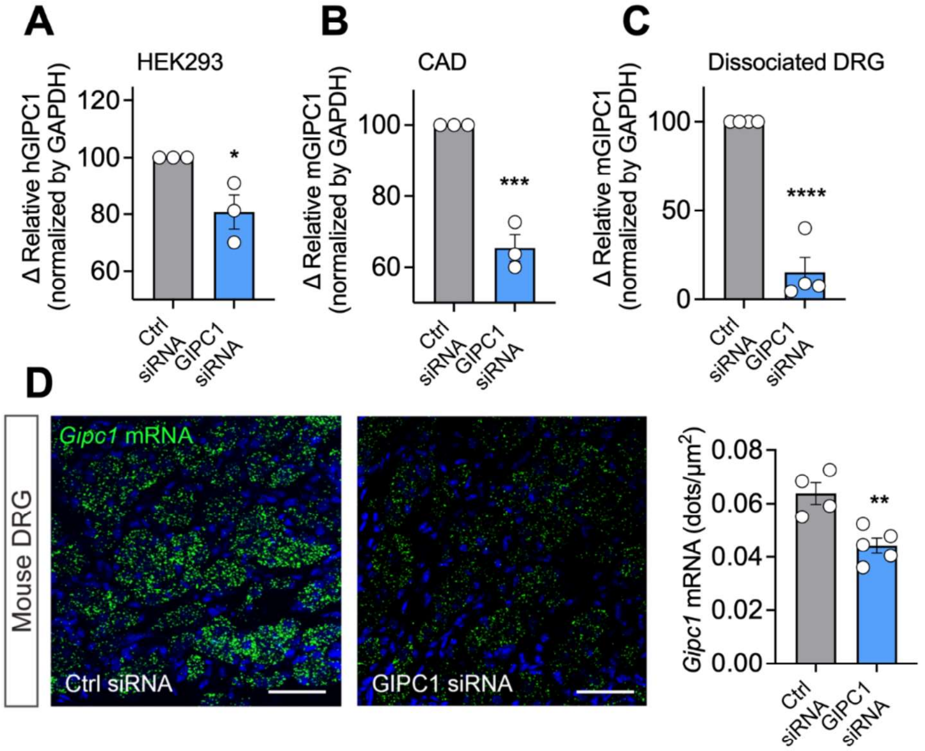

**Figure S6. Confirmation of GIPC1 knockdown using siRNA. A-C.** Quantification of expression of *Gipc1* mRNA in HEK293T (**A**), CAD cells (**B**) or cultured dissociated DRG neurons (**C**) in the presence of control (Ctrl) or GIPC1 siRNA determined by quantitative RT-PCR. Data from 3-4 independent experiments with triplicate wells. **D.** RNAScope$^{TM}$ localization of *Gipc1* mRNA in mouse DRG 48 h after intrathecal injection of Ctrl or GIPC1 siRNA. Representative image from N=5 mice. Mean±SEM. *$P<0.05$, **$P<0.01$, *** $P<0.001$, **** $P<0.0001$. Unpaired t-test.

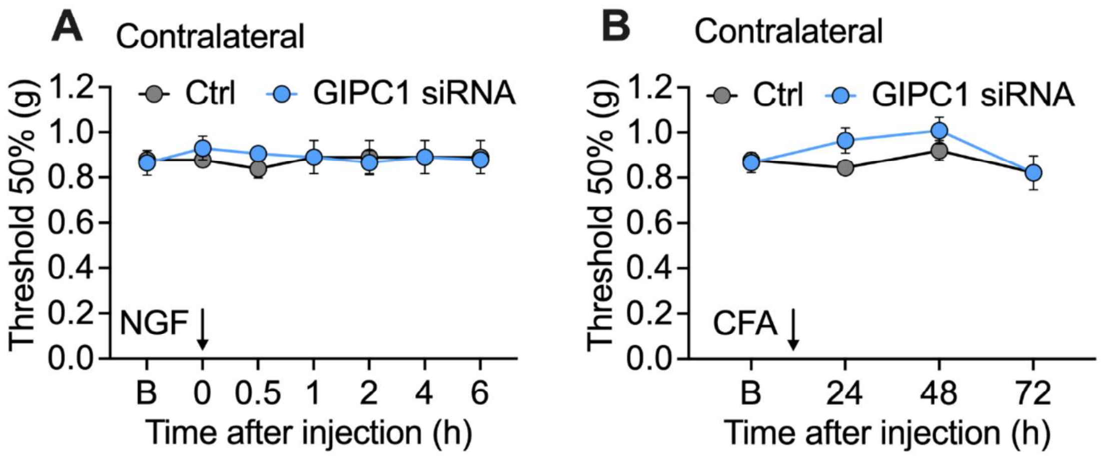

**Figure S7. Withdrawal responses of the contralateral paw.** Withdrawal responses of the contralateral paw to VFF stimulation after injection of NGF (**A**) or CFA (**B**) into the ipsilateral hindpaw. N=6-8 male mice per group.

**SI Tables**

**Table S1. Interfacing residues between NRP1-NGF and NRP1-TrkA in NRP1/NGF/TrkA ternary computational model.**

| NRP1 (monomer 1) | | NGF (monomer 1) | |
|---|---|---|---|
| **Residue** | **H (Hydrogen bond) S (Salt Bridge)** | **Residue** | **H (Hydrogen bond) S (Salt Bridge)** |
| Tyr297 | | His75 | |
| Trp301 | | Trp76 | |
| Thr316 | | Arg114 | H |
| Pro317 | | Ala116 | |
| Gly318 | | Val117 | |
| Glu319 | | Arg118 | HS |
| Asp320 | HS | | |
| Lys351 | | | |
| Tyr353 | H | | |
| Thr413 | H | | |
| Gly414 | | | |
| **NRP1 (monomer 1)** | | **NGF (monomer 2)** | |
| **Residue** | **H (Hydrogen bond) S (Salt Bridge)** | **Residue** | **H (Hydrogen bond) S (Salt Bridge)** |
| Asp320 | | Ser2 | H |
| Thr349 | | Ser3 | |
| Lys351 | | Phe7 | |
| Tyr354 | | His8 | |
| Lys356 | H | Ser73 | |
| Thr357 | | | |
| Thr410 | | | |
| Trp411 | | | |
| Glu412 | | | |
| Thr413 | | | |
| **NRP1 (monomer 1)** | | **TrkA (monomer 1)** | |
| **Residue** | **H (Hydrogen bond) S (Salt Bridge)** | **Residue** | **H (Hydrogen bond) S (Salt Bridge)** |
| Ile107 | | Cys36 | |
| Ala108 | | Pro37 | |
| Pro110 | | Asp38 | |
| Pro201 | | Ser46 | |
| Gly202 | H | Arg52 | |
| Gly203 | | Ala65 | |
| Met204 | | Glu66 | |
| Lys373 | S | Asn67 | H |
| Asn376 | H | Leu68 | |
| Lys377 | S | Thr69 | |
| Pro378 | | Glu92 | |
| | | Arg94 | |
| | | Glu231 | S |
| | | Glu233 | HS |
| | | Gln234 | H |
| | | Asp257 | S |

Interface calculations are made using PISA (12). C-terminal Arg118 from NGF and residues of C-terminal arginine binding pocket from b1 domain of NRP1 are in red. Residues predicted to form or possibly form hydrogen bonds (H) or salt bridges (S) are also indicated

**Table S2. Gating properties of $Ca^{2+}$ and $Na^+$ currents recorded from mouse DRG neurons.**

| **Total $Ca^{2+}$** | **DMSO (0.1%)** | **EG00229 (30 µM)** | **NGF (50 nM)** | **EG00229 + NGF** |
|---|---|---|---|---|
| Activation | | | | |
| $V_{1/2}$ | 5.793 ± 1.891 (10) | -2.828 ± 0.871 (6)† | 5.834 ± 1.995 (9)†† | -0.473 ± 1.265 (8) |
| *k* | 7.756 ± 1.534 (10) | 4.230 ± 0.731 (6) | 7.972 ± 1.608 (9) | 6.428 ± 1.109 (8) |
| Inactivation | | | | |
| $V_{1/2}$ | -27.372 ± 1.919 (9) | -20.137 ± 0.901 (7) | -23.909 ± 3.080 (9) | -20.166 ± 1.314 (8) |
| *k* | -14.944 ± 2.109 (9) | -10.623 ± 0.842 (7) | -15.079 ± 3.305 (9) | -13.176 ± 1.287 (8) |
| **Total $Na^+$** | **DMSO (0.1%)** | **EG00229 (30 µM)** | **NGF (50 nM)** | **EG00229 + NGF** |
| Activation | | | | |
| $V_{1/2}$ | -22.436 ± 1.150 (10) | -23.229 ± 0.499 (13) | -23.151 ± 0.418 (14) | -23.600 ± 0.993 (11) |
| *k* | 5.098 ± 1.014 (10) | 3.580 ± 0.434 (13) | 3.432 ± 0.362 (14) | 5.145 ± 0.877 (11) |
| Inactivation | | | | |
| $V_{1/2}$ | -48.736 ± 4.219 (10) | -43.306 ± 4.249 (13) | -54.450 ± 2.244 (14) | -38.622 ± 4.013 (11)* |
| *k* | -18.568 ± 4.934 (10) | -21.036 ± 5.439 (13) | -14.748 ± 2.243 (14) | -19.732 ± 4.708 (11) |

Values are mean±SEM calculated from fits of the data from the indicated number of individual cells (in parentheses) to the Boltzmann equation; $V_{1/2}$ midpoint potential (mV) for voltage-dependent of activation or inactivation; *k*, slope factor. Data were analyzed with 1-way ANOVA with Tukey's multiple comparisons. † p=0.0115 for $V_{1/2}$ activation of EG00229 vs DMSO (0.1%); †† p=0.0131 for $V_{1/2}$ activation of NGF vs EG00229; * p= 0.0175 for $V_{1/2}$ inactivation of EG00229 + NGF vs DMSO (0.1%).

**Table S3. Pharmacological parameters ($pEC_{50}$) of NGF derived in at human TrkA, in the absence or presence of NRP1 modulation**.

| **CAD cells** | **Control** | **NRP1 inhibition (EG00229)** |
|---|---|---|
| Cytosolic ERK (FRET) | 10.64 ± 0.41 (5) | 10.53 ± 0.17 (5) |
| Nuclear ERK (FRET) | 11.09 ± 0.33 (5) | 10.01 ± 0.38 (5) * |
| **HEK293T cells** | **Control** | **NRP1 overexpression (HaloTag)** |
| Cytosolic ERK (FRET) | 9.89 ± 0.21 (5) | 9.87 ± 0.10 (4) |
| Nuclear ERK (FRET) | 10.37 ± 0.19 (5) | 10.59 ± 0.33 (5) |
| ERK Transcription (SRE-Luc2P) | 9.67 ± 0.48 (8) | 10.15 ± 0.57 (8) *** |
| TrkA-Rluc8 Trafficking (BRET) | 9.06 ± 0.10 (6) | 9.08 ± 0.13 (6) |
| TrkA/TrkA Oligomerization (BRET) | 8.63 ± 0.14 (4) | 8.74 ± 0.09 (5) |

Derived in HEK293T or neuron-like CAD cells transfected with full-length, human TrkA. Values are mean±SEM calculated from non-linear regressions fit of the data from independent replicates (in parentheses), each with triplicate wells. Quantified as the negative antilog of the half-maximal effective concentration ($pEC_{50} = -logEC_{50}$). *$P$<0.05, ***$P$<0.001. Paired t-test.

**Table S4. siRNA sequences used in this study.**

| Target | Sequence |
|---|---|
| Control | UGGUUUACAUGUCGACUAA<br>UGGUUUACAUGUUGUGUGA<br>UGGUUUACAUGUUUUCUGA<br>UGGUUUACAUGUUUUCCUA |
| mNrp1 | GAAUUGCUGUGGAUGAUAU<br>AGUAAGAGGUGUCAUCAUU<br>CCACAAGGUUCAUCAGGAU<br>GGAAUGUUCUGUCGCUAUG |
| hGipc1 | GCACUCGGGCUCACCAUCA<br>GGCCGUACCUUCACGCUGA<br>GCAAGGCCUUCGACAUGAU<br>CUGGAGAGUUACAUGGGUA |
| mGipc1 | GCAUCGAGGGCUUCACUAA<br>CGUCGGCCUUUGAGGAGAA<br>GUGGAUGACUUGCUAGAGA<br>GCUGAGGCCUUCCGACUAC |
